# *Staphylococcus aureus* β-toxin exerts anti-angiogenic effects by inhibiting re-endothelialization and neovessel formation

**DOI:** 10.1101/2021.11.26.470137

**Authors:** Phuong M. Tran, Sharon S. Tang, Wilmara Salgado-Pabón

## Abstract

*Staphylococcus aureus* is the causative agent of numerous severe human infections associated with significant morbidity and mortality worldwide. *S. aureus* often targets the vascular endothelium to interfere with proper host responses during invasive infections. In this study, we provide evidence that *S. aureus* β-toxin inhibits wound repair mechanisms in human endothelial cells by preventing cell proliferation and migration. These findings were confirmed in a rabbit aortic explant model where β-toxin impedes sprout formation. Decreased cell proliferation was accompanied by decreased production of the angiogenic proteins endothelin-1, IGFBP-3, thrombospondin-1, TIMP-1, and TIMP-4. Meanwhile, inhibited wound repair was marked by increased HGF secretion from endothelial cells, likely a marker of endothelial cell damage. Together, these findings establish a mechanistic role for β-toxin where it inhibits proper tissue repair processes that likely promote *S. aureus* infective niche.

## INTRODUCTION

*Staphylococcus aureus* is the causative agent of numerous diseases including skin and soft-tissue infections, bacteremia, toxic shock syndrome, pneumonia, and infective endocarditis (IE) (1). It is also the leading cause of health care-associated infections (2, 3). *S. aureus* facilitates these distinct infections by producing a plethora of secreted and cell-associated virulence factors that, together, enable the organism to bind to, colonize, or invade host cells and tissues, and promote immune system subversion (4–9). The cytolysin β-toxin is encoded by a majority of *S. aureus* strains and is produced during infection by phage excision (10). β-toxin promotes skin and nasal colonization, modulates the immune response to infection, and increases the severity of life-threatening infections like necrotizing pneumonia and IE (10–15). β-toxin exhibits pathogenic properties as a function of its sphingomyelinase (SMase) activity (16, 17). SMases hydrolyze sphingomyelin, a structural molecule in eukaryotic membranes, into phosphocholine and ceramide. Ceramide can be further processed by host enzymes into ceramide-1-phosphate (C1P), sphingosine, and sphingosine-1-phosphate (S1P) (18). These bioactive sphingolipids are widely recognized as essential signaling molecules that regulate various cellular functions and pathological processes, including cell growth and survival, inflammation and immune cell trafficking, vascular integrity and dysfunction, and angiogenesis (19–24).

Angiogenesis, the development of new capillaries from preexisting blood vessels, allows remodeling of the vascular system (25). It requires the coordinated efforts of endothelium-associated cells (e.g. pericytes, fibroblasts, monocytes) to sustain vessel sprouting and for functional maturation and vessel stabilization. Under physiologic conditions, angiogenesis leads to organ growth and re-vascularization of damaged or ischemic tissues for wound healing, while aberrant angiogenesis disrupts these processes and can promote pathological states like malignancy, asthma, diabetes, cirrhosis, multiple sclerosis, and endometriosis (25–27). More recently, angiogenesis induced as a result of microbial infection has been shown to act as an innate immune mechanism to control and clear invading pathogens (28). Not surprisingly, some bacterial pathogens (e.g. *Bartonella* spp., *Mycobacterium tuberculosis* and *Pseudomonas aeruginosa*), viruses (e.g. hepatitis C virus and human papilloma virus), and pathogenic fungi (e.g. *Candida albicans* and *Aspergillus fumigatus*) have also been found to co-opt angiogenic processes to promote disease development and/or persistence (28–31).

*S. aureus* necrotizing pneumonia and IE are prime examples of aggressive infections that present with tissue injury that distinctly lacks signs of healing (14, 32). *S. aureus* IE is characterized by non-healing vegetative lesions, tissue destruction at and around the heart valves, and systemic complications such as ischemic liver lesions or kidney injury (4, 5, 33). Studies have also shown that β-toxin modulates endothelial cell function. In murine pneumonia models, β-toxin induces vascular leakage and neutrophilic inflammation (14). *In vitro*, it increases platelet aggregation and inhibits neutrophil transendothelial migration, processes important in development of IE (12, 15, 32, 34). Furthermore, in human aortic endothelial cells, β-toxin decreases expression of the chemokine IL-8 and upregulates expression of VCAM-1, both of which are important angiogenic molecules (12, 15, 35, 36). Altogether, these studies indicate that a central process may exist where β-toxin targets angiogenesis as a pathogenesis mechanism that enhances *S. aureus* infections.

Therefore, we investigated whether β-toxin modulates the endothelial cell angiogenic response as a possible mechanism for potentiating *S. aureus* infections. We provide evidence that β-toxin specifically targets both human endothelial cell proliferation and cell migration in well-established *in vitro* models. These results are consistent with a dysregulated angiogenic response centered around inhibition of the production of proteins important for these processes. While β-toxin can affect the complexity of capillary-like structures formed *in vitro*, this effect is insufficient to explain β-toxin’s anti-angiogenic properties. Conclusive evidence comes from *ex vivo* studies that demonstrate β-toxin prevents branching microvessel formation, highlighting its ability to interfere with tissue re-vascularization and vascular repair.

## RESULTS

### β-toxin targets the production of angiogenic proteins involved in proliferation and migration

SMase activity results in the production of bioactive sphingolipids, recognized as signaling molecules that modulate angiogenesis (37, 38). Hence, we first sought to establish whether *S. aureus* β-toxin alters the secretion profile of angiogenesis-related proteins in immortalized human aortic endothelial cells (iHAECs). For this purpose, we used a human angiogenesis proteome array to profile 48 proteins in iHAEC supernatants from subconfluent monolayers treated for 24 h under proangiogenic (growth medium ± VEGF), antiangiogenic (+ axitinib), or toxin conditions. In complete medium (basal medium with growth supplements), the most highly detected proteins secreted by endothelial cells were serpin E1, endothelial growth factor (EGF), thrombospondin-1, and endothelin-1, which promote either matrix degradation, growth, or proliferation and migration (Fig. S1A). These were followed largely by proteins that promote endothelial cell proliferation and migration (artemin, insulin-like growth factor binding protein (IGFBP)-2, IGFBP-3, pentraxin 3), capillary formation (angiopoietin-2), and tissue inhibitor of metalloproteinases (TIMP)-1. Hence, as expected, iHAECs in growth medium are under angiogenic-inducing conditions. For the subsequent studies, the secretion profile of iHAECs treated with VEGF (10 ng mL^-1^), axitinib (30 μM), or β-toxin (50 mg mL^-1^) were expressed as fold change from growth medium control, with a cut-off of ≥1.5-fold or ≤0.5-fold as thresholds for 50% increases or decreases in protein levels, respectively (39).

VEGF promotes angiogenesis and is produced by iHAECs during growth in a monolayer, albeit at low levels (Fig. S1A). Thus, VEGF treatment was used to maximally induce angiogenesis under experimental conditions tested in our study. VEGF-treated cells exhibited a similar profile to that of growth medium control, with the exception of IL-8 which showed increased production (1.5-fold; +50%) (Fig. 1). Furthermore, in the presence of complete medium, VEGF did not further induce proliferation, confirming optimal proangiogenic conditions (Fig. S2A, S2B). Axitinib is an antiangiogenic molecule that inhibits VEGF receptor-1, −2, and −3 signaling in endothelial cells (40) and inhibits both cell proliferation and cell migration (41). At working concentrations, axitinib significantly inhibited metabolic activity and proliferation (Fig. S2C, S2D) while preserving integrity of the monolayer as visually established. Consistent with its antiangiogenic activity, axitinib induced an overall inhibitory profile with <0.5-fold decreases in production of granulocyte-macrophage colony-stimulating factor (GM-CSF), A Disintegrin and Metalloproteinase with Thrombospondin Motifs (ADAMTS)-1, Hepatocyte Growth Factor (HGF), pentraxin 3, TIMP-1, and dipeptidyl peptidase IV (DPPIV) (Fig. 1). β-toxin treatment at a sublethal concentration (Fig. S1B) resulted in an overall inhibitory profile that was centered around proteins important for cell proliferation and migration: IGFBP-3 (0.4-fold; −64%), TIMP-1 (0.2-fold; −75%), TIMP-4 (0.5-fold; −50%), thrombospondin-1 (0.4-fold; −63%), and endothelin-1 (0.3-fold; −71%) (Fig.1). The protein profile resulting from β-toxin treatment suggests that β-toxin is an antiangiogenic molecule that inhibits angiogenesis by targeting proliferation and migration.

**Figure 1.**
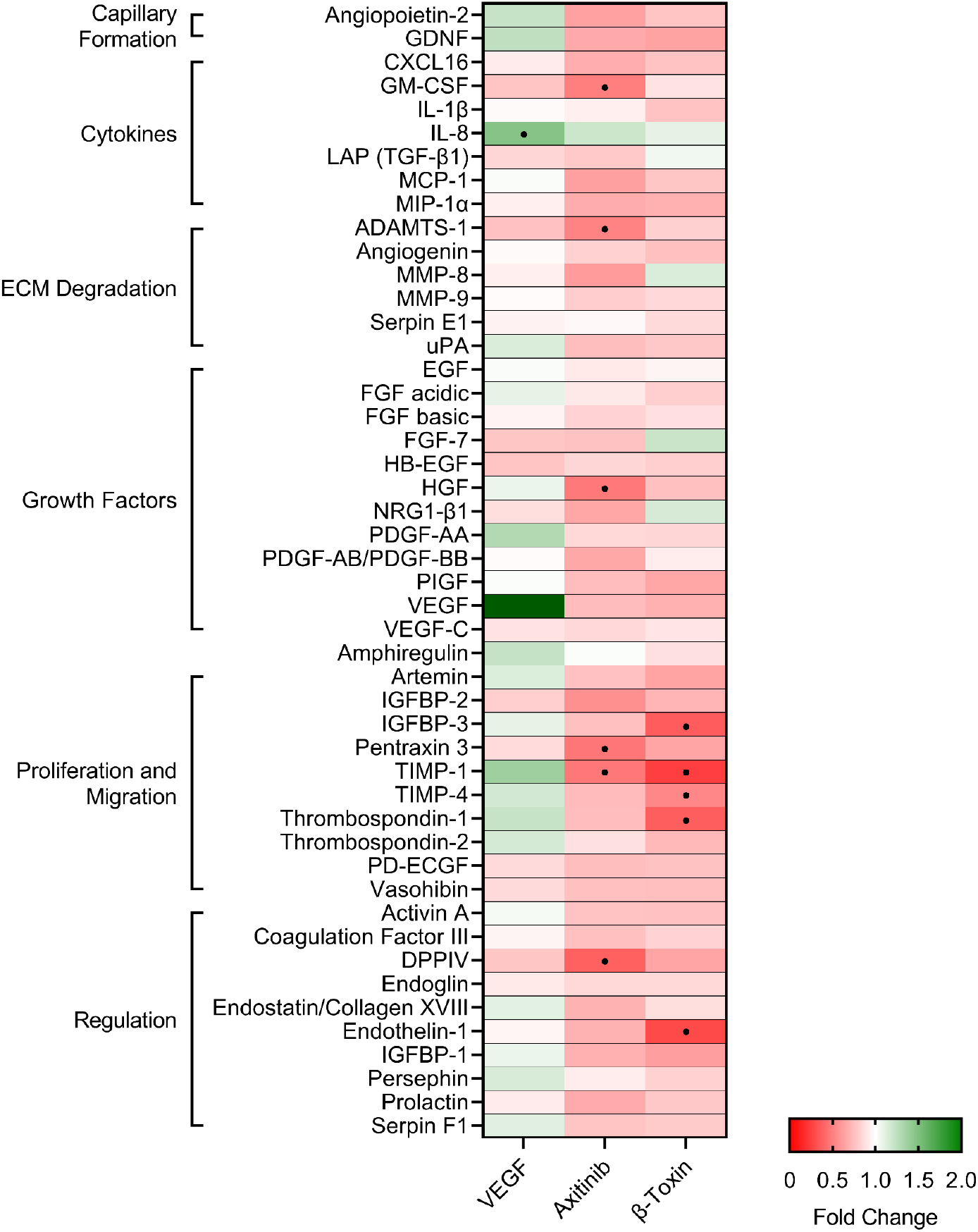
β-toxin inhibits production of angiogenic proteins from human aortic endothelial cells in monolayer growth. Immortalized human aortic endothelial cells (iHAECs) grown to near confluency on gelatin-coated plates were treated with either VEGF (10 ng mL^-1^), axitinib (30 μM), or β-toxin (50 μg mL^-1^) for 24 h. Protein production was assessed by Proteome Profiler Human Angiogenesis Array Kit. Results are the mean fold change over untreated cells of three independent experiments conducted in duplicate. • angiogenic-related factors with a 50% increase (>1.5-fold change) or decrease (<0.5-fold change) from media control.

### β-toxin inhibits wound healing

As *in vitro* wound healing assays are a function of proliferation and migration, we used this approach to directly evaluate the ability of iHAECs to close a gap in the monolayer in the presence or absence of β-toxin (50 μg mL^-1^). VEGF (10 ng mL^-1^) and axitinib (10 μM) were used as inducing or inhibition controls, respectively. Of note, axitinib, at the concentration used to treat monolayers (30 μM) resulted in generalized iHAEC toxicity during wound healing (as observed by loss of integrity of the monoloayer) (Fig. S2B). Hence, we decreased the axitinib concentration to 10 μM for this assay which inhibited proliferation but did not result in cytotoxicity (Fig. S3).

Without treatment, iHAECs closed 80% of the gap by 24 h, as measured in time-lapse analyses (Fig. 2). iHAECs stimulated with VEGF showed an increase in percent gap closure of 15%, while those treated with axitinib displayed significant inhibition with a 28% decrease in gap closure compared to untreated cells (Fig. 2A and 2B). Similarly, iHAECs treated with β-toxin exhibited a 28.5% decrease in gap closure (Fig. 2A and 2C). During infection, host cells can undergo prolonged exposure to β-toxin before vascular damage occurs, potentiating the inhibitory phenotype. To assess this, iHAECs were treated overnight with β-toxin, prior to gap formation, and thereafter as previously performed. Pretreatment with β-toxin significantly inhibited gap closure by 34.3% (Fig. 2A and 2D), but overall provided no significant further inhibition (Fig. 2B and 2C). These results were confirmed in human umbilical vein endothelial cells (HUVECs), where similar inhibition of gap closure was observed upon exposure to β-toxin with and without pretreatment (Fig. S4). These data indicate that β-toxin markedly inhibits re-endothelialization, contributing to defects in vascular repair.

**Figure 2.**
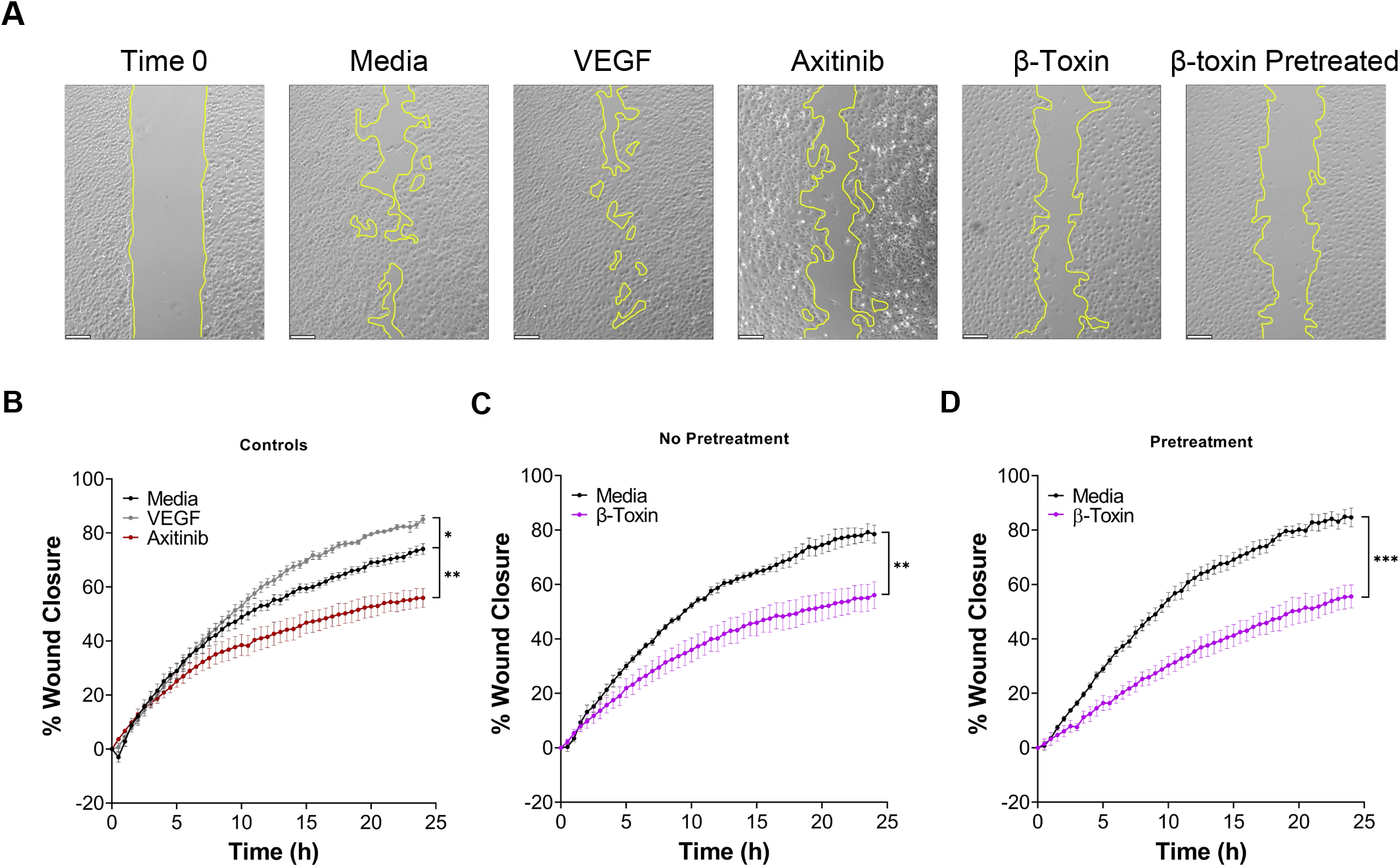
β-Toxin inhibits wound healing. Time course analysis of iHAECs grown to confluency in silicone inserts that create uniform gaps upon removal and treated with either VEGF (10 ng mL^-1^), axitinib (10 μM), or β-toxin (50 μg mL^-1^) for 24 h. Images captured every 30 min. (A) Phase-contrast microscopy at Time 0 (representative image) and at 24 h for all conditions tested. Scale bar = 200 μm. (B) iHAECs treated with VEGF or axitinib. (C) iHAECs treated with β-toxin. (D) iHAECs pretreated overnight with β-toxin prior to gap formation and thereafter. (B-D) All results are mean ± SEM of five independent experiments with four replicates each. * *p* < 0.0332, ** *p* < 0.0021, *** *p* < 0.002, \*\*\*\**p* = <0.0001; two-way repeated measures ANOVA with Tukey’s multiple comparisons test.

### Differential protein production during wound healing

Because differential effects were observed in the wound healing assay in response to β-toxin and experimental controls, we investigated the proteome profile in media collected at the end of the assays. This provided an opportunity to establish whether distinct profiles would ensue from cell populations that contain cells actively migrating and proliferating to close a gap versus those in a monolayer. In the wound healing assay, serpin E1, EGF, and endothelin-1 remained unchanged compared to growth in a monolayer (Fig. S1A). Overall, these remained the most highly detected proteins secreted by iHAECs *in vitro*. Interestingly, compared to growth in a monolayer, iHAECs in the wound healing assay exhibited both increases and decreases in proteins involved in growth, proliferation, and migration, where placental growth factor (PlGF), TIMP-1, and thrombospondin-1 were increased by 1.6-fold (+57%), 2.1-fold (+108%), and 1.7-fold (+67%), respectively, while hepatocyte growth factor (HGF), and IGFBP-2 were decreased by 0.5-fold (−51%) and 0.4-fold (−57%), respectively (Fig. S1A).

iHAECs in the wound healing assay were more responsive to the effects of VEGF and axitinib. Cells treated with VEGF exhibited an induced angiogenesis profile compared to both untreated cells (Fig. 3) and VEGF-treated cells grown in a monolayer (Fig. S5). Proteins that showed increases during wound healing spanned most categories, with those important in extracellular matrix degradation being the sole exception (Fig. 3). These results are consistent with enhanced gap closure (Fig. 2A) and reiterate the prominent effects of VEGF on regulation of angiogenic factors. While axitinib significantly inhibited gap closure (Fig. 2A), it induced a mixed protein profile and a shift away from the inhibitory profile observed in iHAEC monolayers (Fig. S5). Instead, during wound healing, axitinib specifically targeted matrix metalloproteinase (MMP)-8 (0.5-fold; −47%), PDGF-AA (1.7-fold; +34%), and TIMP-4 (0.5-fold; −41%) (Fig. 3). Subtle decreases in a select group of proteins caught our attention as they stand opposite to the changes observed in VEGF-treated cells. These included the cytokines CXCL-16 and IL-1β, MMP-9, the growth factor HB-EGF, and proteins involved in regulating or aiding proliferation and migration (IGFBP-2, TIMP-1, activin A, serpin F1, and DPPIV).

**Figure 3.**
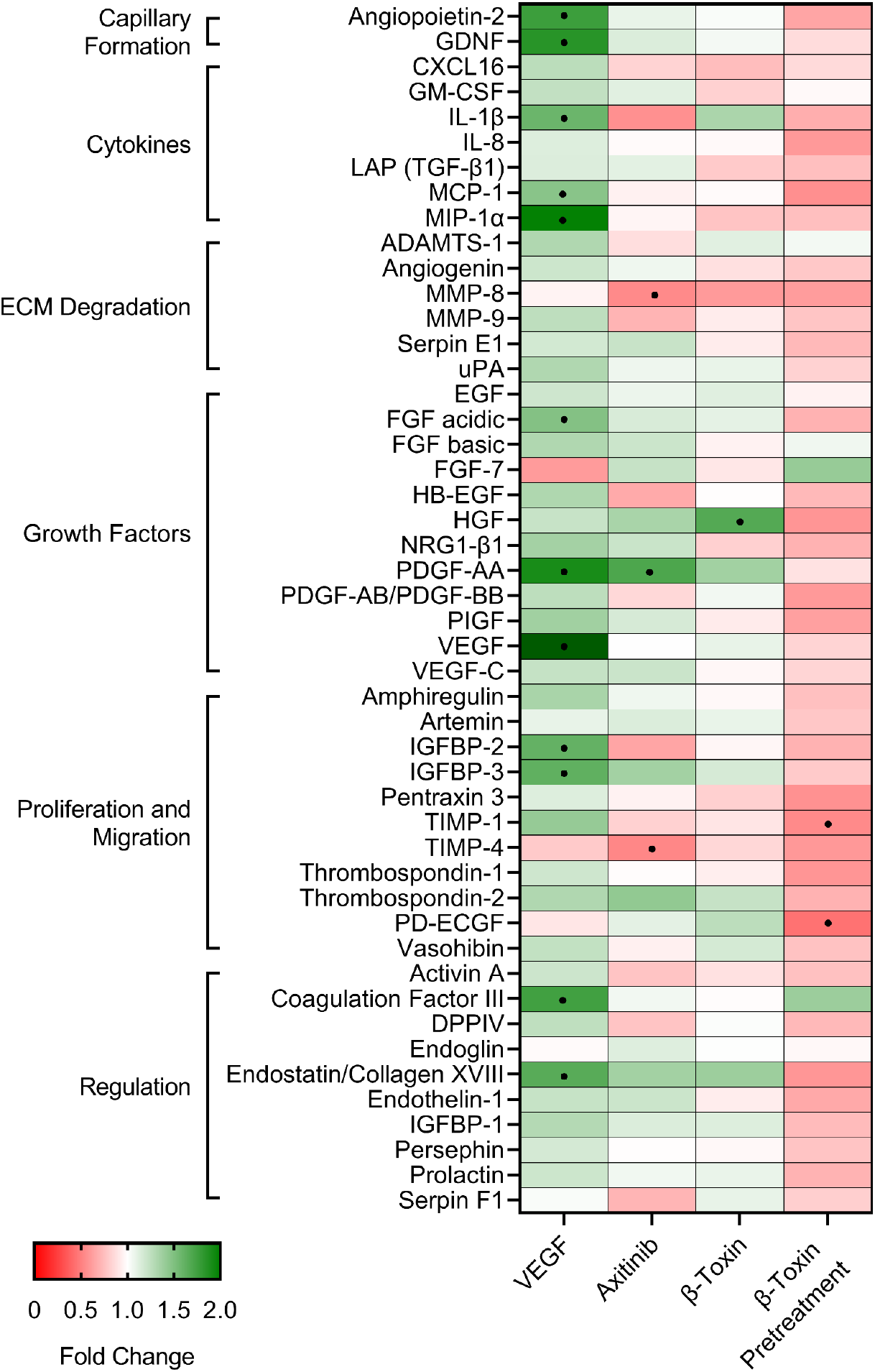
β-Toxin modulates production of angiogenic proteins from iHAECs during wound healing. iHAECs grown to confluency in silicone inserts that create uniform gaps upon removal and treated with either VEGF (10 ng mL^-1^), axitinib (10 μM), or β-toxin (50 μg mL^-1^) for 24 h. Angiogenesis proteome arrays were determined from culture supernatants collected at 24 h. Results shown are the mean fold change from untreated cells of five independent experiments. • angiogenic-related factors with a 50% increase (>1.5-fold change) or decrease (<0.5-fold change) from media control.

Strikingly, iHAECs treated with β-toxin during wound healing exhibited a mostly neutral profile with subtle changes observed throughout. Most notably were decreased MMP-8 (important in matrix remodeling for angiogenesis) and increased endostatin (an angiogenesis inhibitor) (Fig. 3). The most relevant change after β-toxin treatment was increased production of the growth factor HGF by 1.7-fold (+69%) (Fig. 3). This profile is intriguing given that in that same context β-toxin significantly inhibited gap closure (Fig. 2C). Pretreatment shifted the profile back to mostly inhibitory, where iHAECs exhibited large decreases in proliferation/migration proteins PD-ECGF (0.4-fold; −59%) and TIMP-1 (0.5-fold; −46%) (Fig. 3). Therefore, subtle changes in the balance of angiogenic proteins may be sufficient to significantly impact wound healing as measured *in vitro*. Collectively, these results illustrate the context–dependent characteristics of the angiogenic profile of iHAECs in response to exogenous agents.

### β-toxin inhibits cell migration and cell proliferation

Having established that β-toxin inhibits wound healing, we addressed whether this effect was the result of impaired cell migration, cell proliferation, or both. For this purpose, we conducted wound healing assays in the presence of mitomycin C, an antiproliferative agent. Mitomycin C was used at 2 μg mL^-1^ as this concentration reduced metabolic activity by 49.2%, and in combination with β-toxin did not further decrease metabolic activity (Fig. 4A). Treatment with mitomycin C above 2 μg mL^-1^ resulted in cell toxicity (data not shown). Mitomycin C inhibited wound healing by 17.8% compared to untreated cells, where the largest effect on cell proliferation and concomitant decreases in wound closure were measured past 15 h (Fig. 4B, 4C). iHAECs concurrently treated with β-toxin (50 μg mL^-1^) and mitomycin C exhibited a 40.9% decrease in wound closure compared to untreated cells and a 28.8% decrease compared to those treated only with mitomycin C (Fig. 4B and 4C). The additive effect of mitomycin C and β-toxin in the wound healing assay is consistent with β-toxin inhibiting cell migration.

**Figure 4.**
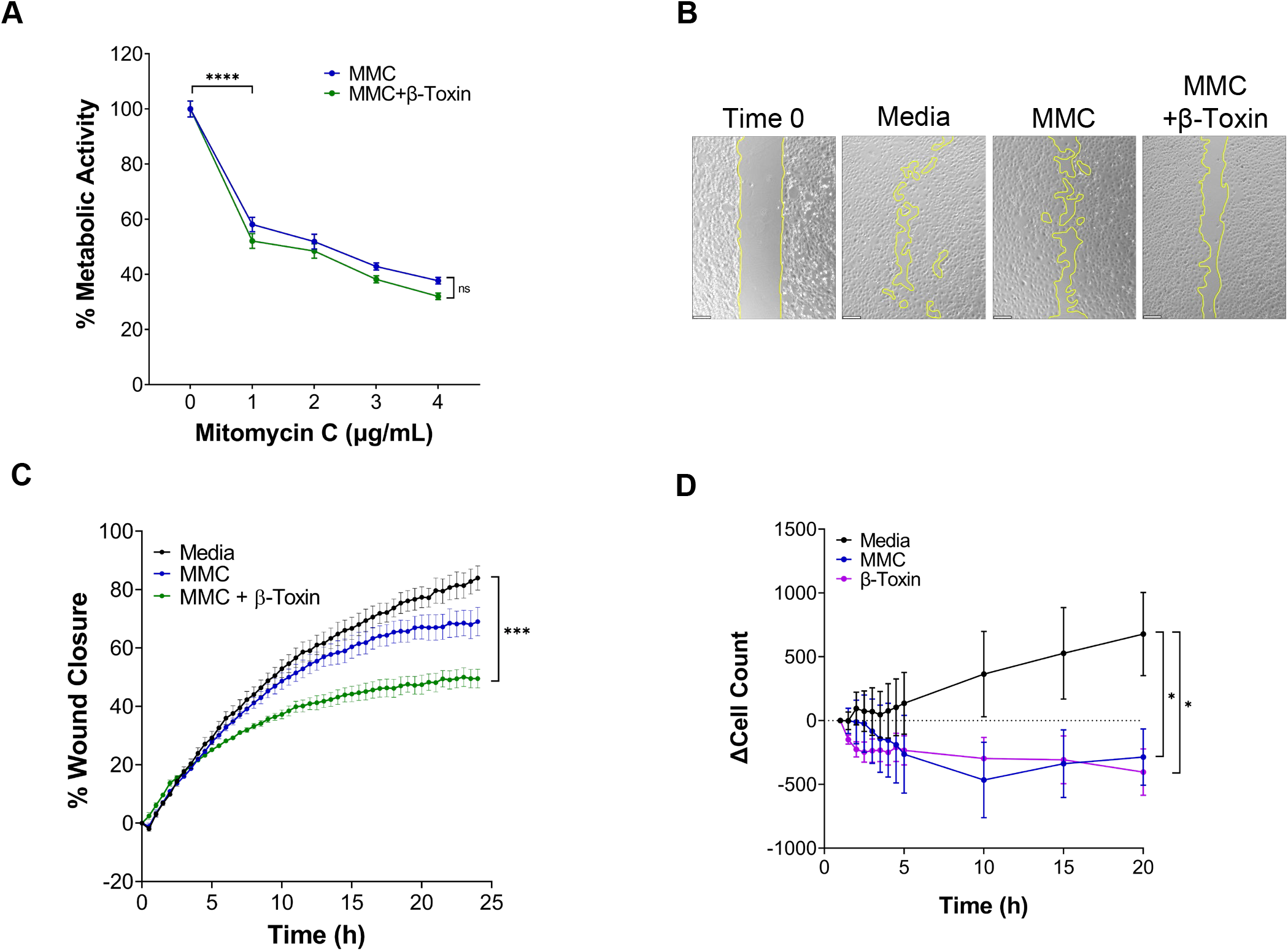
β-toxin inhibits migration and proliferation. (A) Percent metabolic activity. iHAECs grown to near confluency on 1% gelatin-coated plates and treated for 24 h with mitomycin C (MMC) in the absence or presence of β-toxin (50 μg mL^-1^). **** *p* = <0.0001; two-way repeated measures ANOVA. Unpaired, two-tailed t test was used to compare individual MMC treatments to media control (0 μg mL^-1^). (B) Phase-contrast microscopy at Time 0 (representative image) and at 24 h for all conditions tested. Images captured every 30 min. Scale bar = 200 μm. (C) Percent wound closure over time of iHAECs treated with MMC (2 μg mL^-1^) ± β-toxin (50 μg mL^-1^). All results are mean ± SEM of five independent experiments with four replicates each. \*\*\**p* < 0.002; two-way repeated measures ANOVA. (D) Cell proliferation of iHAECs seeded at 7,000 cells/well and treated with MMC (2 μg mL^-1^) or β-toxin (50 μg mL^-1^) over a 20-h period. Cells counted every 30 min for the first 5 h then every 5 h thereafter. Results represent the change in cell count (mean ± SEM) of three independent experiments conducted in triplicate. * *p* < 0.0332, unpaired, two-tailed t-test at 20 h.

In the wound healing assay, initially cells exist in a monolayer and gap closure is largely driven by actively migrating cells of the gap cell front. Therefore, we sought to directly test β-toxin effects on cell proliferation in a population of actively proliferating cells. For this, iHAECs were cultured at the time of seeding in the presence or absence of β-toxin or mitomycin C and total cell counts were measured from images taken at various time points over a 20-h period (Fig. 4D). In this context, β-toxin significantly inhibited cell proliferation comparable to the inhibition induced by mitomycin C (Fig. 4D). These data provide evidence that β-toxin can interfere with the angiogenic process by both inhibiting cell migration and cell proliferation.

### β-toxin inhibits neovessel formation in rabbit aortic ring explants

Re-vascularization is the ultimate outcome of angiogenesis. *In vitro*, endothelial cell differentiation into capillaries or tubulogenesis can be addressed with a tube formation assay. For this assay, iHAECs and HUVECs were seeded on growth factor reduced (GFR)-Matrigel in medium containing serum to induce tube formation. Cells were treated with either β-toxin or axitinib (inhibitor control) at the time of seeding and the average tube length and loop count was measured over a 12-h time frame. Axitinib significantly decreased tube length and loop count over time (Fig. 5A-C). Yet, β-toxin treatment produced mixed results in aortic versus umbilical vein endothelial cells. In the presence of β-toxin, the average tube length and loop count remained unaffected in iHAECs (Fig. 5D, 5E). With HUVECs, β-toxin had no effect on tube length but did significantly decrease loop count, suggesting a direct effect on network complexity during re-vascularization (Fig. 5F, 5G). The differential responses of HUVECs and iHAECs to β-toxin likely reflects either the heterogeneity of endothelial cells or the altered physiology of immortalized cells (42).

**Figure 5.**
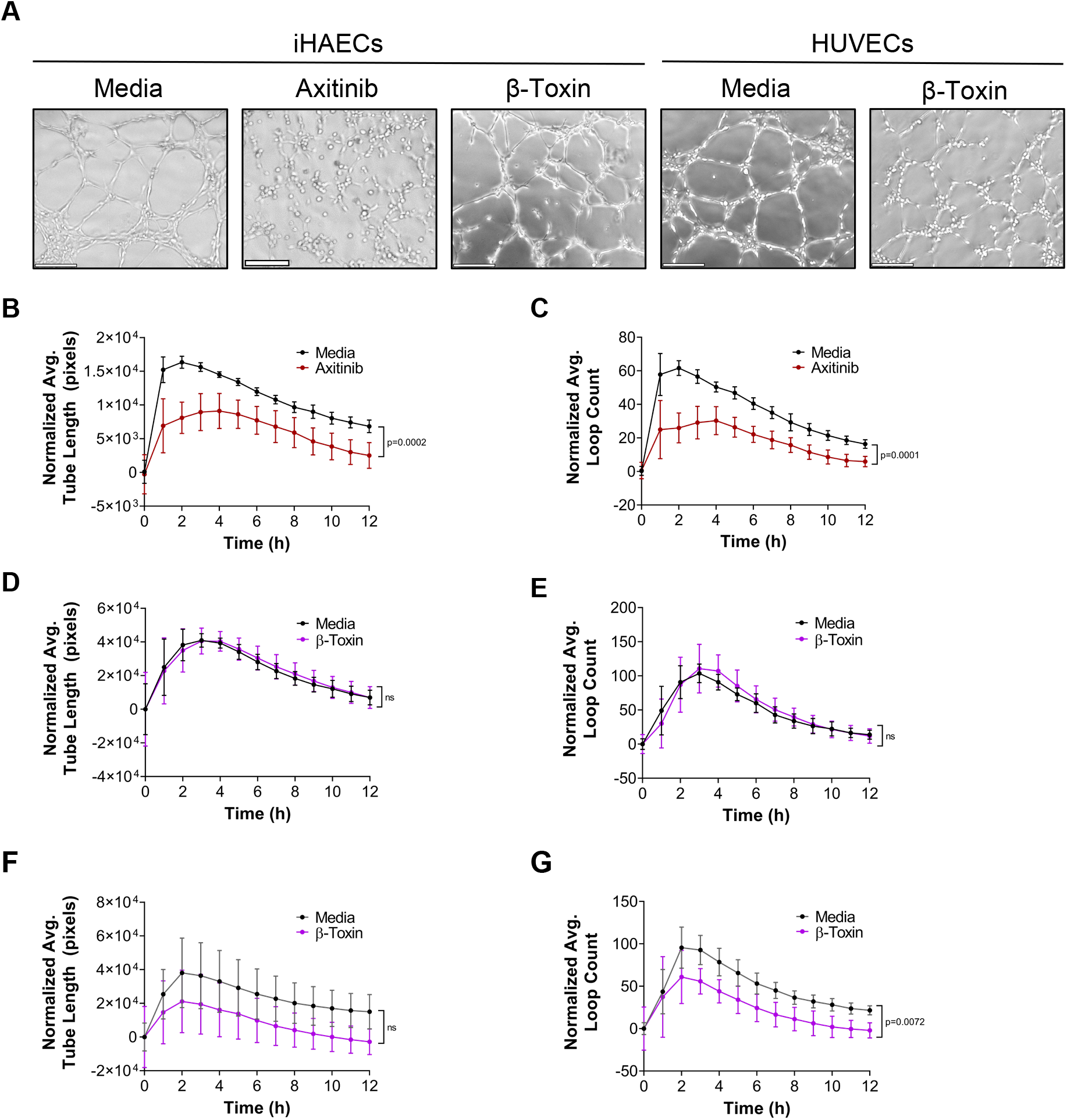
β-Toxin has differential effects on tube formation. iHAECs seeded on GFR-Matrigel were treated with either axitinib (30 μM) or β-toxin (50 μg mL^-1^) and tube formation imaged every 1 h for 12 h. (A) Phase-contrast microscopy at 3 h. Scale bar = 200 μm. (B) Average tube length over time of iHAECs ± axitinib. (C) Average loop count over time of iHAECs ± axitinib. (D) Average tube length over time of iHAECs ± β-toxin. (E) Average loop count over time of iHAECs ± β-toxin. (F) Average tube length over time of HUVECs ± β-toxin. (G) Average loop count over time of HUVECs ± β-toxin. (B-G) Results are means ± SD for at least 6 independent experiments with five replicates each. Statistical significance determined by two-way repeated measures ANOVA.

Hence, it remained to be established if β-toxin inhibits angiogenesis by targeting capillary formation. To address this in a more physiologically relevant system, we utilized the rabbit aortic ring model of angiogenesis. In this model, thoracic and abdominal aortas were explanted from New Zealand white rabbits, cut into ~1 mm sections, embedded into a thin layer of GFR-Matrigel, and cultured in complete medium to induce sprouting at the severed edge of the explant. We used equal numbers of thoracic and abdominal aortic rings per condition obtained from 3 individual rabbits. After embedding, aortic rings were cultured in the presence or absence of β-toxin or axitinib for 14 days. Untreated aortic ring explants (n=26) formed sprouts within a week that continued to grow in density and complexity over time while rings treated with axitinib (n=8) or β-toxin (n=12) failed to sprout over the course of the experiment (Fig. 6; Fig. S6; Fig. S7). Thus, in the more physiologically relevant context of the aortic ring model, β-toxin completely inhibited angiogenesis.

**Figure 6.**
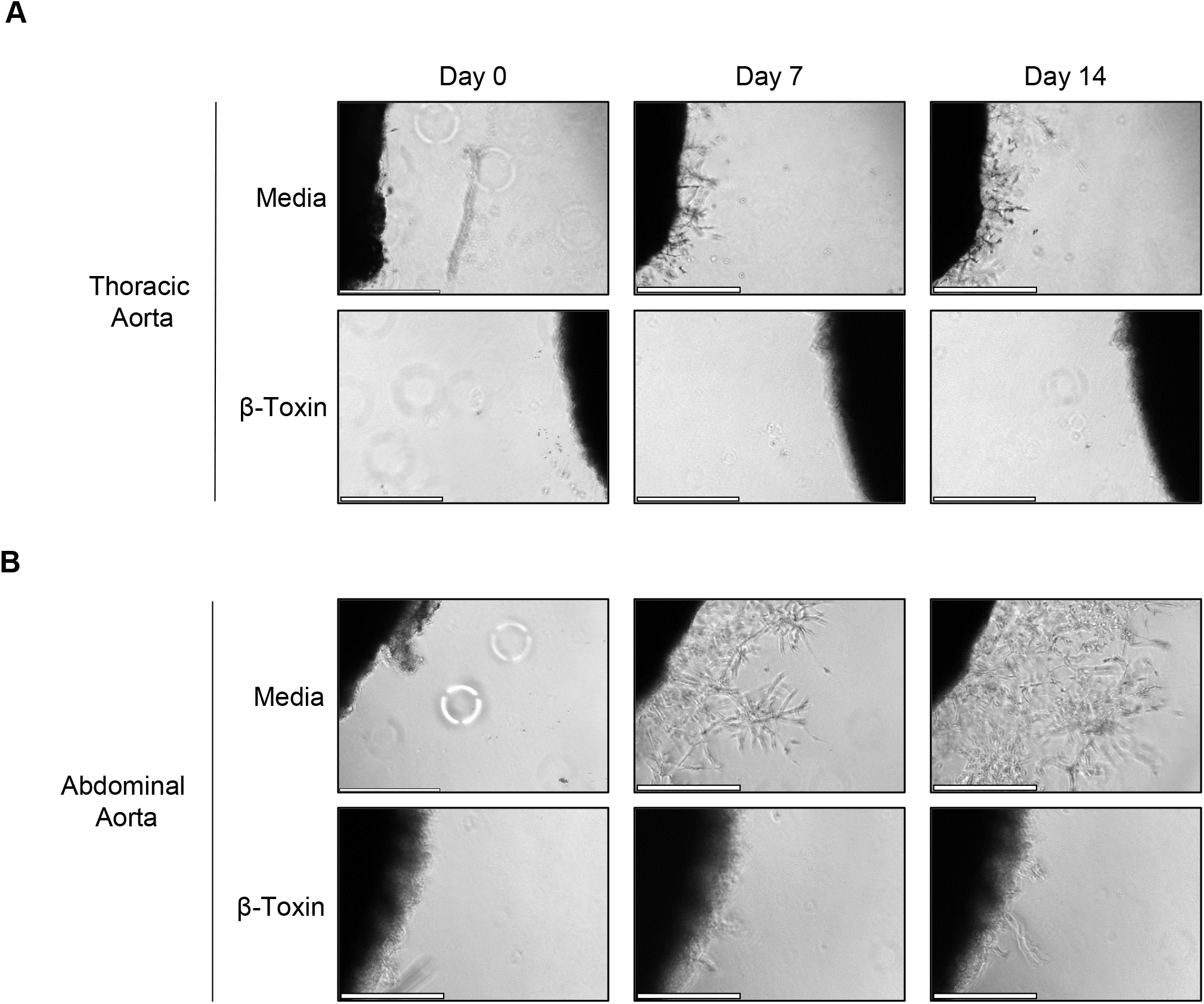
β-toxin inhibits sprout formation. Thoracic and abdominal aortas were collected and sectioned from 2–3 kg New Zealand white rabbits. Rings were cultured on GFR-Matrigel in the presence or absence of β-toxin (50 μg mL^-1^). Scale bar = 500 μm. (A) Phase-contrast microscopy of thoracic aortic rings. (B) Phase-contrast microscopy of abdominal aortic rings.

## DISCUSSION

The vascular endothelium reaches into every organ, where endothelial cells, as building blocks of the vascular network, maintain cardiovascular homeostasis and health of surrounding tissue (43, 44). Angiogenesis, the process of developing neovessels from pre-existing formations, is a critical function of the endothelium and essential for vascular repair and tissue re-vascularization after injury. Hence, given the contribution of β-toxin to the exacerbation of *S. aureus* pneumonia (10, 14) and vegetative lesions during IE (10, 45, 46), we addressed whether β-toxin interferes with angiogenic processes. With the use of the *ex vivo* rabbit aortic ring model, which preserves the microenvironment of the aortic endothelium, we demonstrate that β-toxin is an anti-angiogenic virulence factor that prevents branching microvessel formation. We provide evidence that β-toxin specifically targets both human endothelial cell proliferation and cell migration as tested *in vitro*, which is consistent with an angiogenic response dysregulated in the production of proteins important in cell proliferation and cell migration. Furthermore, while β-toxin can affect the complexity of capillary-like structures formed *in vitro*, this effect is insufficient to explain β-toxin’s anti-angiogenic properties. Yet, the different sensitivities of iHAECs and HUVECs to the effect of β-toxin in the tube formation assay suggest that β-toxin could induce increased pathology on tissues where endothelial cells are more sensitive to the toxin. These results highlight a mechanism where β-toxin exacerbates *S. aureus* invasive infections by interfering with tissue re-vascularization and vascular repair.

Ischemic or injured tissues release factors into the environment to trigger sprouting angiogenesis (47, 48). The balance between stimulatory (pro-angiogenic) and inhibitory (anti-angiogenic) factors controls the angiogenic switch, where endothelial cells change from a quiescent to a sprouting phenotype (49). When the local concentration of angiogenic inducers is produced in excess of the angiogenic inhibitors, neovessel formation is triggered. The angiogenic switch is off when the local concentration of angiogenic inhibitors overpowers the stimulators. Growth factors that promote angiogenesis include VEGF, HGF, serpin E1, EGF, bFGF, endothelin-1, and PDGF, while those that turn it off include thrombospondin-1 and endostatin (48, 50). Aberrant angiogenesis occurs when the system is inappropriately or chronically activated, or when there is a spatiotemporal imbalance of pro- and anti-angiogenic factors. Improper angiogenesis can lead to endothelial dysfunction, malignancy, insufficient wound healing, and various diseases such as retinopathies, fibrosis, diabetes, cirrhosis, and endometriosis (25, 51–53).

*In vitro*, endothelial cell monolayers under pro-angiogenic conditions produce an angiogenesis proteome profile consistent with cells that are triggered to sprout. VEGF stimulation under this condition only further promotes angiogenesis by inducing the release of IL-8, confirming the pro-angiogenic state of endothelial cells. β-toxin shifted the overall profile, where the abundance of many proteins was decreased. β-toxin decreased the production of both endothelin-1 and thrombosponin-1. These two proteins are some of the most highly expressed in iHAECs in our experimental conditions and have opposing effects on angiogenesis. Endothelin-1 is a potent endothelial cell mitogen shown to stimulate migration and to contribute to endothelial cell integrity, of particular importance in newly formed blood vessels (54). Thrombosponin-1 is a non-structural extracellular matrix protein and a potent endogenous inhibitor of cell adhesion, migration, and proliferation (55). Its primary function is to counter the effect of angiogenic stimuli, effectively turning the angiogenic switch off. Thrombosponin-1 decreases may potentially correspond to concomitant decreases of endothelin-1 and/or overall decreases of angiogenic signals. β-toxin also decreased the production of TIMP-1, TIMP-4, and IGFBP-3. TIMPs are known to control MMPs activities to maintain extracellular matrix homeostasis while promoting sprout formation, vessel stabilization, and vessel pruning (regression) (56). TIMPs also possess several cellular functions independent of their MMP-inhibiting activities (57). For example, TIMP-1 promotes cell growth and limits cell migration by controlling focal adhesions (58). TIMPs functions are spatiotemporally regulated, and dysregulation causes a functional imbalance leading to excessive and uncontrolled matrix degradation resulting in sprouting defects, vessel instability and/or vascular regression (56, 57). IGFBP-3 is yet another multifunctional protein with context-dependent effects on angiogenesis. In HUVECs, IGFBP-3 disrupts established focal adhesions and actin stress fibers inhibiting cell migration, while in endothelial progenitor cells, it stimulates cell proliferation, migration, and survival to promote vascular repair (59, 60). Therefore, the cell type dictates whether IGFBP-3 induces or inhibits cell migration. Altogether, these results indicate that β-toxin likely causes an imbalance in protein production that cumulatively disrupts angiogenesis in iHAECs.

The wound healing assay provided a context with which to address the effect of β-toxin on the endothelium’s endogenous capacity to repair. It mimics re-endothelialization after vascular injury, a process dependent on cell migration and proliferation. iHAECs were sensitive to VEGF stimulation in this context, producing an array of growth factors known to promote angiogenesis (PDGF, FGF, and ANG-2), but in particular, factors that induce vessel maturation and capillary network formation (ANG-2 and GDNF) (61, 62). In this context, the IGFBPs and TIMP-1 are increased as well as coagulation factor III (also known as tissue factor) and several cytokines. The angiogenesis proteome profile is consistent with cells that are not only triggered to sprout but also ready for re-vascularization and tissue repair. During wound healing, β-toxin largely induced production of HGF, with subtle increases in PDGF-AA and endostatin, and a subtle decrease in MMP-8. Excess HGF in the serum is clinically used as an indicator of advanced atherosclerotic lesions, vascular lesions, and hypertension (63–65). Vascular lesions are accompanied by endothelial cell injury. As such, it has been suggested that endothelial cells produce HGF to promote repair of damaged endothelial cells at these lesions (63–65). Hence, iHAECs might induce HGF as a protective mechanism in response to endothelial injury caused by β-toxin. This response is consistent with increases in PDGF-AA, an early factor produced by senescent endothelial cells at cutaneous wound sites that promotes tissue repair (66). MMP-8 is a pro-angiogenic factor rapidly induced during tissue injury that stimulates proliferation, migration, and capillary network formation (67). Therefore, the decrease in MMP-8 in combination with an increase in endostatin (angiogenic inhibitor), while subtle, may be relevant in the context of wound healing. Alternatively, the inhibitory effect of β-toxin in wound healing could be directly driven by sphingolipid metabolites produced from SMase activity.

The sphingolipid metabolites ceramide and S1P are critical regulators of cellular and pathological processes yet have opposing effects on vascular functions (23, 24, 37, 68). Ceramide is the first sphingolipid metabolite produced from β-toxin’s hydrolysis of sphingomyelin. It promotes cellular functions associated with endothelial dysfunction and inhibition of angiogenesis. Ceramide is a well-known antiproliferative molecule and induces endothelial barrier dysfunction, oxidative stress, cell senescence, and cell death. It inhibits cell migration by disassembling focal adhesions and depolymerizing stress fibers (69). Ceramide can further be metabolized into S1P. S1P promotes cellular functions associated with maintenance of vascular integrity and induction of angiogenesis. It stimulates cell proliferation, supports barrier integrity, and promotes cell survival. S1P enhances cell contacts with the extracellular matrix to induce cell migration (70, 71). At the end, the cellular balance between ceramide and S1P dictates the outcomes. This balance is also known as the ceramide rheostat (24). Interestingly, several proteins regulated by β-toxin (IGFBP-3, TIMP-1, thrombospondin-1, HGF, endothelin-1) are either controlled by ceramide/S1P or regulate their activity. IGFBP-3 activates sphingosine kinase to convert sphingosine into S1P stimulating growth and promoting cell survival (72). Furthermore, IGFBP-3 directly inhibits sphingomyelinase (73). Meanwhile, ceramide has been shown to downregulate TIMP-1 in human glioma cells resulting in reduced tumor volume (74). Exogenous C2-ceramide causes apoptosis of porcine thyroid cells by decreasing thrombospondin-1 expression (75). Conversely, S1P induces TIMP-1 production (76). HGF is protective against ceramide-mediated apoptosis (77) while endothelin-1 induces SMase activity resulting in increased VCAM-1 surface expression. The fate of ceramide following production by β-toxin is not known. Altogether, the anti-angiogenic effects of β-toxin described herein are consistent with ceramide rheostat signaling. Future studies will be directed at elucidating the underlying cellular processes driving the anti-angiogenic endothelial cell phenotype in the presence of β-toxin. In particular, the physiological context and sphingolipid metabolites that mediates those responses.

Angiogenesis is a highly complex but fundamental physiological process essential for vascular injury repair (i.e., due to mechanical damage or toxin-mediated damage of the endothelium) as well as the re-vascularization of ischemic or injured tissue (i.e., due to embolic events, trauma, or caused by pathogens and their toxins). Here, we provide evidence that *S. aureus* β-toxin inhibits capillary formation by a mechanism that targets cell proliferation and cell migration. β-toxin inhibition of IGFBP-3 and TIMP-1 are of particular interest as these molecules play crucial roles in endothelial cell proliferation and migration and are linked to SMase activity. While it is not clear how sphingolipid metabolites inhibit the abundance of IGFBP-3, decreases in IGFBP-3 favors ceramide accumulation as opposed to the more protective sphingolipid S1P (73). Ceramide not only arrests cell growth but also regulates production of TIMP-1 (19, 20, 74). During wound healing, β-toxin can also target MMP-8 to limit endothelial cell proliferation and migration, while turning angiogenesis off by increasing the levels of endostatin. In conclusion, β-toxin is an anti-angiogenic virulence factor that can prevent proper vascular repair, keeping the endothelium in a proinflammatory, hypercoagulable state, and preventing neovessel formation. This environment in turn would allow *S. aureus* to maintain its infectious niche.

## ACKNOWLEDGEMENTS

This work was supported by NIH grant 1R01AI134692-01A1 to W.S.-P.

## AUTHOR CONTRIBUTIONS

Conceptualization, P.M.T and W.S.-P.; Methodology, P.M.T and W.S.-P.; Formal Analysis, P.M.T; Investigation, P.M.T and S.T.; Resources, W.S.-P.; Writing – original draft, P.M.T and W.S.-P.; Writing – review & editing, P.M.T, S.T., and W.S.-P.; Visualization, P.M.T.; Supervision, W.S.-P.; Funding Acquisition, W.S.-P.

## DECLARTION OF INTERESTS

The authors declare no competing interests.

## MATERIALS AND METHODS

### Protein Expression and Purification

N-terminal His6-tagged β-toxin was previously cloned into *E. coli* TOP10 using a pTrcHis TOPO vector (45). The plasmid was maintained with 100 μg mL^-1^ carbenicillin in all growths. Cells were grown in 1 L terrific broth (24 g yeast extract, 12 g tryptone, 4 mL glycerol, 100 mL of supplement [0.17 M KH_2_PO_4_, 0.72 M K_2_HPO_4_]) at 37°C to an OD_600_ 0.4 – 0.8 followed by induction with 1 mM IPTG overnight at 30°C. Pelleted cells were resuspended in 25 mL resuspension buffer (50 mM NaH_2_PO_4_, 500 mM NaCl, 20 mM imidazole, pH 8) and three Pierce Protease Inhibitor Mini Tablets. 10 mL aliquots were divided into bead lysing tubes containing 7 g of 0.1 mm glass beads and homogenized using a PreCellys Cryolys Evolution (bertin Instruments) with the following settings: 9900 rpm, 6-30 s cycles with 60 s rests, 4°C. Lysate was centrifuged (40 min, 50,000 x g, 4°C) and clarified with a 0.45 μm filter. Affinity chromatography with HisPur™ Cobalt Resin followed by an imidazole-gradient elution (50 mM NaH_2_PO_4_, 500 mM NaCl, 250 mM imidazole, pH 8) was used to separate β-toxin. Protein-containing fractions, assessed by SDS-PAGE, were dialyzed against 4 L PBS, pH 7.4, overnight at 4°C. Protein concentration was determined by Qubit™ using the Qubit™ Protein Broad Range kit. Typical yield was 4-5 mg per liter of growth. Purity was assessed by Coomassie stain of SDS-PAGE gels and was at least 95% by visual observation. All proteins were cleaned of endotoxin via Detoxi-Gel™ resin and endotoxin levels were assessed using the ToxinSensor™ LAL Endotoxin Assay Kit. Proteins were used when the final endotoxin concentration in experiments was ≤0.025 ng mL^-1^ (78).

### Culture Conditions

Immortalized human aortic endothelial cells (iHAECs) are a recently established cell line shown to retain phenotypic and functional characteristics of primary cells, serving as a large-vessel model system in which to address questions relevant to vascular biology (78). Human umbilical vein endothelial cells (HUVECs) were obtained from Thermo Fisher as low passage cells.

Cells were grown at 37°C, 5% CO_2_ in phenol red–free, endothelial cell basal medium (Medium 200) supplemented with low-serum growth supplement (LSGS, final concentrations of: FBS 2%, hydrocortisone 1μg mL^-1^, human epidermal growth factor 10 ng mL^-1^, basic fibroblast growth factor, 3 ng mL^-1^, heparin 10 μg mL^-1^). Cells were maintained on 1% gelatin-coated plates unless otherwise stated. Cells were passaged at least twice before use in experiments. iHAECs were used at passages between 4 and 10 from a single clone. Primary HUVECs were used between 4 and 12 passages. Mycoplasma-testing was conducted every 6 months using MycoAlert™ Plus Mycoplasma Detection Kit.

### Cell Growth and Metabolic Activity

An MTS assay was used to determine cell viability. Cells were seeded at 7,000 cells/well into gelatin-coated 96-well plates and grown overnight to near confluency. Media was removed and replaced with 100 μL of media containing increasing concentrations of β-toxin, VEGF, axitinib, or mitomycin C followed by overnight incubation. 20 μL of CellTiter 96® AQueous One Solution was added to each well followed by a 1 h incubation at 37°C, 5% CO_2_. A plate reader was used to read absorbance at 490 nm. Three independent experiments in triplicate were conducted. The data were normalized so that untreated cells were 100% activity.

### Proteome Profiler™ Human Angiogenesis array

Gelatin-coated 96-well tissue culture plates were seeded at 7,000 cells/well and grown to near confluence. Fresh media containing β-toxin at 50 μg mL^-1^ was added, and plates were incubated for 24 h at 37°C, 5% CO_2_. The conditioned media was removed and stored at −80°C until analyzed. Each treatment was conducted in triplicate with three technical replicates. The relative expression of 55 angiogenesis-related proteins was determined from the conditioned media of various experiments using a Proteome Profiler™ Human Angiogenesis Antibody Array according to the manufacturer’s instructions modified for fluorescent analysis. 120 μL of conditioned media was incubated with a cocktail of biotinylated detection antibodies for 1 h at room temperature. During this incubation, the membrane containing the capture antibodies was blocked at room temperature. After the hour incubation, the sample-antibody mixture was added to the washed membrane and incubated overnight at 4°C. After a series of washes, the membrane was incubated with IRDye 800CW Streptavidin (1:2000 dilution) for 30 min at room temperature in the dark. After a series of washes, the fluorescent signal was detected using an Azure c600 (Azure Biosystems; 120 μm resolution, auto intensity). The signal produced at each spot is proportional to the amount of analyte bound and the mean pixel intensity of the duplicate spots on the membrane was calculated and averaged using Image Studio Software (LI-COR). Fold-changes over untreated controls were calculated for each detected protein. All treatments were matched. After an extensive literature search and cross-referencing of the GTExPortal and Expression Atlas databases, seven analytes unlikely to be produced by endothelial cells were removed from final analysis (angiopoietin-1, angiostatin/plasminogen, EG-VEGF, FGF-4, leptin, platelet factor 4, serpin B5). None of these analytes were produced by iHAECs.

### Wound Healing Assay

4-chamber silicone inserts were placed into 12-well uncoated tissue culture treated plates. Each chamber was seeded with 3.08×10^4^ cells and plates were incubated at 37°C, 5% CO_2_ for 4 h. The media was removed and replaced media containing β-toxin (50 μg mL^-1^) and incubated overnight. The inserts were removed, and conditioned media was saved. The wells were washed with DPBS, and the conditioned media was returned to the wells with additional media containing effectors so that the final volume was 1.5 mL per well. Experiments were also conducted where β-toxin (50 μg mL^-1^), VEGF (10 ng mL^-1^), axitinib (10 μM), or mitomycin C (2 μg mL^-1^) was added after insert removal. The plates were incubated overnight in a Leica DMi8 equipped with a Tokai Hit stage-top incubator set to 37°C, 5% CO_2_. Images were captured every 30 min for 24 h using a HC PL FLUOTAR 4x/0.13 objective lens. Five independent experiments were conducted for each treatment condition. Images were automatically analyzed via ImageJ. The edges were found (*Process* → *Find Edges*) and the image was smoothed 10 times (*Process* → *Smooth*). A MinError(I) threshold was then applied (*Image* → *Adjust* → *Auto Local Threshold: MinError (I)*) to automatically detect cells. The particle count was then quantified with a particle size of 1000-infinity (*Analyze* → *Analyze Particles [size: 1,000 – infinity]*) (79).

### Cell Proliferation Assay

For cell proliferation assays, cells were seeded at 7,000 cells/well into gelatin-coated 96-well plates and immediately treated with either β-toxin (50 μg mL^-1^), VEGF (10 ng mL^-1^), axitinib (10 μM), or mitomycin C (2 μg mL^-1^). Plates were incubated overnight in a Leica DMi8 equipped with a Tokai Hit stage-top incubator (Tokai Hit Co., Ltd.) set to 37°C, 5% CO_2_. Merged images were captured every 30 min for 5 h, then every 5 h using a HC PL FLUOTAR 4x/0.13 objective lens. Three independent experiments were conducted for each treatment condition. Cells were automatically counted using ImageJ by modifying the protocol outlined by Venter and Niesler (2019). Images were converted to grayscale and the edges found (*Process* → *Find Edges*). An Isodata threshold was then applied (*Image* → *Adjust* → *Auto Threshold: Isodata*) to automatically detect cells. The particle count was then quantified after determining the appropriate particle size to decrease background (*Analyze* → *Analyze Particles [size: 0.003 – 0.2]*).

### Tube Formation

Wells in angiogenesis μ-slides were coated with 10 μL of GFR-Matrigel and allowed to polymerize for 1 h in a humidified chamber at 37°C, 5% CO_2_. Cells were seeded at 10,000 cells/well in media containing β-toxin (50 μg mL^-1^) or axitinib (30 μM). The μ-slides were incubated overnight in a Leica DMi8 equipped with a Tokai Hit stage-top incubator set to 37°C, 5% CO_2_. Images were captured every hour for 12 h using a HC PL FLUOTAR 4x/0.13 objective lens. Images were analyzed via the ImageJ Angiogenesis Analyzer plugin (80). A minimum of six independent experiments with five technical replicates were conducted.

### Aortic Ring Explant

Mixed-sex New Zealand white rabbits, 2-3 kg, were purchased from Charles River and maintained at Charmany Instructional Facility at the School of Veterinary Medicine (SVM) of the University of Wisconsin (UW)-Madison. All rabbits were individually caged and given access to food and water *ad libitum*. Rabbits were given a period of at least four days to acclimate and deemed generally healthy by a veterinarian before experimental procedures. All experimental procedures were conducted under a UW-Madison SVM IACUC-approved protocol (#V006222).

Aortic ring explants were conducted by modifying the thin-layer method (81). The thoracic and abdominal aortas were excised immediately after euthanasia. In a petri dish containing PBS, excess fascia and connective tissue were removed, then 1 – 1.5 mm^2^ cross-sections were cut with a scalpel. 300 uL phenol red-free GFR-Matrigel was added to wells in 24-well plates and rings immediately embedded. After 10 min polymerization at 37°C, 5% CO_2_, 500 μL supplemented Medium 200 was added and plates were incubated at 37°C, 5% CO_2_ up to 14 days. Medium 200 contained LSGS, 100 U mL^-1^ penicillin-streptomycin, 2.5 μg mL^-1^ amphotericin B, and relevant treatments. Media (± treatments) was changed every 3-5 days. Merged images were captured every other day using the Leica DMi8 equipped with a Tokai Hit stage-top incubator set to 37°C, 5% CO_2_ using a HC PL FLUOTAR 4x/0.13 objective lens. Growth was assessed using ImageJ (82). A total of three rabbits were used with a minimum of three rings per condition.

### Quantification and Statistical Analysis

Statistical analyses were performed using GraphPad Prism software. For each experiment, the precision measures and number of technical and biological replicates are indicated in figure legends. The number of cells, number of measurements and timing of experiments can be found in the Method Details for each experimental setup. For the proteomics analysis, emphasis was placed on proteins with mean fold change outside of the 0.5-to-1.5-fold change previously described using this same array (39). Cell proliferation, tube formation, and wound healing were analyzed by two-way repeated measures ANOVA with α = 0.05. For MTS assays unpaired two-tailed t-tests were conducted. Statistical significance was given as * *p* < 0.0332, ** *p* < 0.0021, *** *p* < 0.002, **** *p* < 0.0001.

## REFERENCES

1. Salgado-Pabón W, Schlievert PM. 2014. Models matter: the search for an effective *Staphylococcus aureus* vaccine. Nat Rev Microbiol 12:585–591.

2. Wisplinghoff H, Bischoff T, Tallent SM, Seifert H, Wenzel RP, Edmond MB. 2003. Nosocomial bloodstream infections in US hospitals: Analysis of 24,179 cases from a prospective nationwide surveillance study. Clin Infect Dis 39:309–317.

3. Klein E, Smith DL, Laxminarayan R. 2007. Hospitalizations and deaths caused by methicillin-resistant *Staphylococcus aureus*, United States, 1999–2005. Emerg Infect Dis 13:1840–1846.

4. King JM, Kulhankova K, Stach CS, Vu BG, Salgado-Pabón W. 2016. Phenotypes and virulence among *Staphylococcus aureus* USA100, USA200, USA300, USA400, and USA600 clonal lineages. mSphere 1:e00071–16.

5. Spaulding AR, Salgado-Pabón W, Kohler PL, Horswill AR, Leung DYM, Schlievert PM. 2013. Staphylococcal and Streptococcal Superantigen Exotoxins. Clin Microbiol Rev 26:422–447.

6. Dumont AL, Torres VJ. 2014. Cell Targeting by the *Staphylococcus aureus* Pore-Forming Toxins: It’s Not Just About Lipids. Trends Microbiol 22:21–27.

7. Bien J, Sokolova O, Bozko P. 2011. Characterization of Virulence Factors of *Staphylococcus aureus*: Novel Function of Known Virulence Factors That Are Implicated in Activation of Airway Epithelial Proinflammatory Response. J Pathog 2011.

8. Oliveira D, Borges A, Simões M. 2018. *Staphylococcus aureus* Toxins and Their Molecular Activity in Infectious Diseases. Toxins (Basel) 10.

9. Brown AF, Leech JM, Rogers TR, McLoughlin RM. 2014. *Staphylococcus aureus* Colonization: Modulation of Host Immune Response and Impact on Human Vaccine Design. Front Immunol 4.

10. Salgado-Pabón W, Herrera A, Vu BG, Stach CS, Merriman JA, Spaulding AR, Schlievert PM. 2014. *Staphylococcus aureus* β-toxin production is common in strains with the β-toxin gene inactivated by bacteriophage. J Infect Dis 210:784–792.

11. Verkaik NJ, Benard M, Boelens HA, de Vogel CP, Nouwen JL, Verbrugh HA, Melles DC, van Belkum A, van Wamel WJB. 2010. Immune evasion cluster-positive bacteriophages are highly prevalent among human *Staphylococcus aureus* strains, but they are not essential in the first stages of nasal colonization. Clin Microbiol Infect 17:343–348.

12. Tajima A, Iwase T, Shinji H, Seki K, Mizunoe Y. 2009. Inhibition of endothelial interleukin-8 production and neutrophil transmigration by *Staphylococcus aureus* beta-hemolysin. Infect Immun 77:327–334.

13. Katayama Y, Baba T, Sekine M, Fukuda M, Hiramatsu K. 2013. Beta-Hemolysin Promotes Skin Colonization by *Staphylococcus aureus*. J Bacteriol 195:1194–1203.

14. Hayashida A, Bartlett AH, Foster TJ, Park PW. 2009. *Staphylococcus aureus* beta-toxin induces lung injury through syndecan-1. Am J Pathol 174:509–518.

15. Herrera A, Kulhankova K, Sonkar VK, Dayal S, Klingelhutz AJ, Salgado-Pabón W, Schlievert PM. 2017. Staphylococcal β-toxin modulates human aortic endothelial cell and platelet function through sphingomyelinase and biofilm ligase activities. MBio 8:e00273–17.

16. Doery HM, Magnusson BJ, Gulasekharam J, Pearson JE. 1965. The Properties of Phospholipase Enzymes in Staphylococcal Toxins. J Gen Microbiol 40:283–296.

17. Doery HM, Magnusson BJ, Cheyne IM, Gulasekharam J. 1963. A Phospholipase in Staphylococcal Toxin which Hydrolyses Sphingomyelin. Nature 198:1091–1092.

18. Chalfant CE, Spiegel S. 2005. Sphingosine-1-phosphate and ceramide 1-phosphate: Expanding roles in cell signaling. J Cell Sci 118:4605–4612.

19. Schenck M, Carpinteiro A, Grassmé H, Lang F, Gulbins E. 2007. Ceramide: Physiological and pathophysiological aspects. Arch Biochem Biophys 462:171–175.

20. Gulbins E, Pin LL. 2006. Physiological and pathophysiological aspects of ceramide. Am J Physiol - Regul Integr Comp Physiol 290:R11–26.

21. Woodcock J. 2006. Sphingosine and Ceramide Signalling in Apoptosis. IUBMB Life 58:462–466.

22. Chatterjee S. 1998. Sphingolipids in atherosclerosis and vascular biology. Arterioscler Thromb Vasc Biol 18:1523–1533.

23. Hait NC, Maiti A. 2017. The Role of Sphingosine-1-Phosphate and Ceramide-1-Phosphate in Inflammation and Cancer. Mediators Inflamm 2017:4806541.

24. Cogolludo A, Villamor E, Perez-Vizcaino F, Moreno L. 2019. Ceramide and regulation of vascular tone. Int J Mol Sci 20.

25. Carmeliet P. 2005. Angiogenesis in life, disease and medicine. Nature 438:932–936.

26. Hsu T, Nguyen-Tran H-H, Trojanowska M. 2019. Active roles of dysfunctional vascular endothelium in fibrosis and cancer. J Biomed Sci 26.

27. Bluff JE, Brown NJ, Reed MW, Staton CA. 2008. Tissue factor, angiogenesis and tumour progression. Breast Cancer Res 10:204.

28. Rolando M, Buchrieser C. 2019. A Comprehensive Review on the Manipulation of the Sphingolipid Pathway by Pathogenic Bacteria. Front Cell Dev Biol 0:168.

29. Tsukamoto K, Shinzawa N, Kawai A, Suzuki M, Kidoya H, Takakura N, Yamaguchi H, Kameyama T, Inagaki H, Kurahashi H, Horiguchi Y, Doi Y. 2020. The *Bartonella* autotransporter BafA activates the host VEGF pathway to drive angiogenesis. Nat Commun 11.

30. Rogers MS, Christensen KA, Birsner AE, Short SM, Wigelsworth DJ, Collier RJ, D’Amato RJ. 2007. Mutant Anthrax Toxin B Moiety (Protective Antigen) Inhibits Angiogenesis and Tumor Growth. Cancer Res 67:9980–9985.

31. Vasil ML, Stonehouse MJ, Vasil AI, Wadsworth SJ, Goldfine H, Bolcome III RE, Chan J. 2009. A Complex Extracellular Sphingomyelinase of *Pseudomonas aeruginosa* Inhibits Angiogenesis by Selective Cytotoxicity to Endothelial Cells. PLoS Pathog 5.

32. Liesman RM, Pritt BS, Maleszewski JJ, Patel R. 2017. Laboratory diagnosis of infective endocarditis. J Clin Microbiol 55:2599–2608.

33. Kinney KJ, Tran PM, Gibson-Corley KN, Forsythe AN, Kulhankova K, Salgado-Pabón W. 2019. Staphylococcal Enterotoxin C promotes *Staphylococcus aureus* Infective Endocarditis Independent of Superantigen Activity. bioRxiv doi.org/10.1101/2019.12.13.875633.

34. Chen Y, Ye LJ, Wu Y, Shen BZ, Zhang F, Qu Q, Qu J. 2020. Neutrophil-lymphocyte ratio in predicting infective endocarditis: A case-control retrospective study. Mediators Inflamm 2020.

35. Kong D-H, Kim YK, Kim MR, Jang JH, Lee S. 2018. Emerging Roles of Vascular Cell Adhesion Molecule-1 (VCAM-1) in Immunological Disorders and Cancer. Int J Mol Sci 19.

36. Heidemann J, Ogawa H, Dwinell MB, Rafiee P, Maaser C, Gockel HR, Otterson MF, Ota DM, Lugering N, Domschke W, Binion DG. 2003. Angiogenic effects of interleukin 8 (CXCL8) in human intestinal microvascular endothelial cells are mediated by CXCR2. J Biol Chem 278:8508–8515.

37. Van Brocklyn JR, Williams JB. 2012. The control of the balance between ceramide and sphingosine-1-phosphate by sphingosine kinase: Oxidative stress and the seesaw of cell survival and death. Comp Biochem Physiol Part B 163:26–36.

38. Mehra VC, Jackson E, Zhang XM, Jiang X-C, Dobrucki LW, Yu J, Bernatchez P, Sinusas AJ, Shulman GI, Sessa WC, Yarovinsky TO, Bender JR. 2014. Ceramide-activated phosphatase mediates fatty acid-induced endothelial VEGF resistance and impaired angiogenesis. Am J Pathol 184:1562–1576.

39. Barron GA, Goua M, Wahle KWJ, Bermano G. 2017. Circulating levels of angiogenesis-related growth factors in breast cancer: A study to profile proteins responsible for tubule formation. Oncol Rep 38:1886–1894.

40. Kelly RJ, Rixe O. 2009. Axitinib-a selective inhibitor of the vascular endothelial growth factor (VEGF) receptor. Target Oncol 4:297–305.

41. Siedlecki J, Wertheimer C, Wolf A, Liegl R, Priglinger C, Priglinger S, Eibl-Lindner K. 2017. Combined VEGF and PDGF inhibition for neovascular AMD: anti-angiogenic properties of axitinib on human endothelial cells and pericytes in vitro. Graefe’s Arch Clin Exp Ophthalmol 255:963–972.

42. Ribatti D, Nico B, Vacca A, Roncali L, Dammacco F. 2002. Endothelial cell heterogeneity and organ specificity. J Hematother Stem Cell Res 11:81–90.

43. Rajendran P, Rengarajan T, Thangavel J, Nishigaki Y, Sakthisekaran D, Sethi G, Nishigaki I. 2013. The Vascular Endothelium and Human Diseases. Int J Biol Sci 9:1057.

44. Félétou M. 2011. Multiple Functions of the Endothelial CellsThe Endothelium: Part 1: Multiple Functions of the Endothelial Cells—Focus on Endothelium-Derived Vasoactive Mediators. Morgan & Claypool Life Sciences, San Rafael.

45. Herrera A, Vu BG, Stach CS, Merriman JA, Horswill AR, Salgado-Pabón W, Schlievert PM. 2016. *Staphylococcus aureus* β-Toxin mutants are defective in biofilm ligase and sphingomyelinase activity, and causation of infective endocarditis and sepsis. Biochemistry 55:2510–2517.

46. Huseby MJ, Kruse AC, Digre J, Kohler PL, Vocke JA, Mann EE, Bayles KW, Bohach GA, Schlievert PM, Ohlendorf DH, Earhart CA. 2010. Beta toxin catalyzes formation of nucleoprotein matrix in staphylococcal biofilms. Proc Natl Acad Sci 107:14407–14412.

47. Adair TH, Montani J-P. 2010. Overview of AngiogenesisAngiogenesis. Morgan & Claypool Life Sciences, San Rafael.

48. Moccia F, Negri S, Shekha M, Faris P, Guerra G. 2019. Endothelial Ca2+ Signaling, Angiogenesis and Vasculogenesis: Just What It Takes to Make a Blood Vessel. Int J Mol Sci 20.

49. Baeriswyl V, Christofori G. 2009. The angiogenic switch in carcinogenesis. Semin Cancer Biol 19:329–337.

50. Bussolino F, Di Renzo MF, Ziche M, Bocchietto E, Olivero M, Naldini L, Gaudino G, Tamagnone L, Coffer A, Comoglio PM. 1992. Hepatocyte growth factor is a potent angiogenic factor which stimulates endothelial cell motility and growth. J Cell Biol 119:629–641.

51. Johnson DE, O’Keefe RA, Grandis JR. 2018. Targeting the IL-6/JAK/STAT3 signalling axis in cancer. Nat Rev Clin Oncol 15:234–248.

52. Iredale JP, Pellicoro A, Fallowfield JA. 2017. Liver Fibrosis: Understanding the Dynamics of Bidirectional Wound Repair to Inform the Design of Markers and Therapies. Dig Dis 35:310–313.

53. Karska-Basta I, Pociej-Marciak W, Chrząszcz M, Kubicka-Trząska A, Dębicka-Kumela M, Gawęcki M, Romanowska-Dixon B, Sanak M. 2021. Imbalance in the Levels of Angiogenic Factors in Patients with Acute and Chronic Central Serous Chorioretinopathy. J Clin Med 10:1–15.

54. Dong F, Zhang X, Wold LE, Ren Q, Zhang Z, Ren J. 2005. Endothelin-1 enhances oxidative stress, cell proliferation and reduces apoptosis in human umbilical vein endothelial cells: role of ETB receptor, NADPH oxidase and caveolin-1. Br J Pharmacol 145:323–333.

55. Lawler J. 2002. Thrombospondin-1 as an endogenous inhibitor of angiogenesis and tumor growth. J Cell Mol Med 6.

56. Cabral-Pacheco GA, Garza-Veloz I, la Rosa CC-D, Ramirez-Acuña JM, Perez-Romero BA, Guerrero-Rodriguez JF, Martinez-Avila N, Martinez-Fierro ML. 2020. The Roles of Matrix Metalloproteinases and Their Inhibitors in Human Diseases. Int J Mol Sci 21:9739.

57. Grünwald B, Schoeps B, Krüger A. 2019. Recognizing the Molecular Multifunctionality and Interactome of TIMP-1. Trends Cell Biol 29:6–19.

58. Akahane T, Akahane M, Shah A, Connor CM, Thorgeirsson UP. 2004. TIMP-1 inhibits microvascular endothelial cell migration by MMP-dependent and MMP-independent mechanisms. Exp Cell Res 301:158–167.

59. Kim J-H, Choi DS, Lee O-H, Oh S-H, Lippman SM, Lee H-Y. 2011. Antiangiogenic antitumor activities of IGFBP-3 are mediated by IGF-independent suppression of Erk1/2 activation and Egr-1-mediated transcriptional events. Blood 118:2622–2631.

60. Lee H-J, Lee J-S, Hwang SJ, Lee H-Y. 2015. Insulin-like growth factor binding protein-3 inhibits cell adhesion via suppression of integrin β4 expression. Oncotarget 6:15163.

61. Akwii RG, Sajib MS, Zahra FT, Mikelis CM. 2019. Role of Angiopoietin-2 in Vascular Physiology and Pathophysiology. Cells 8:471.

62. Zhong Z, Gu H, Peng J, Wang W, Johnstone BH, March KL, Farlow MR, Du Y. 2016. GDNF secreted from adipose-derived stem cells stimulates VEGF-independent angiogenesis. Oncotarget 7:36829–36841.

63. Nishimura M, Ushiyama M, Ohtsuka K, Nishida M, Inoue N, Matsumuro A, Mineo T, Yoshimura M. 1999. Serum Hepatocyte Growth Factor as a Possible Indicator of Vascular Lesions. J Clin Endocrinol Metab 84:2475–2480.

64. Javadi J, Heidari-Hamedani G, Schmalzl A, Szatmári T, Metintas M, Aspenström P, Hjerpe A, Dobra K. 2021. Syndecan-1 Overexpressing Mesothelioma Cells Inhibit Proliferation, Wound Healing, and Tube Formation of Endothelial Cells. Cancers (Basel) 13.

65. Nakamura Y, Morishita R, Nakamura S, Aoki M, Moriguchi A, Matsumoto K, Nakamura T, Higaki J, Ogihara T. 1996. A Vascular Modulator, Hepatocyte Growth Factor, Is Associated With Systolic Pressure. Hypertension 28:409–413.

66. Demaria M, Ohtani N, Youssef SA, Rodier F, Toussaint W, Mitchell JR, Laberge R-M, Vijg J, Van Steeg H, Dollé MET, Hoeijmakers JHJ, de Bruin A, Hara E, Campisi J. 2014. An essential role for senescent cells in optimal wound healing through secretion of PDGF-AA. Dev Cell 31:722–733.

67. Fang C, Wen G, Zhang L, Lin L, Moore A, Wu S, Ye S, Xiao Q. 2013. An important role of matrix metalloproteinase-8 in angiogenesis in vitro and in vivo. Cardiovasc Res 99:146–155.

68. Shea BS, Tager AM. 2012. Sphingolipid Regulation of Tissue Fibrosis. Open Rheumatol J 6:123–129.

69. Presa N, Gomez-Larrauri A, Dominguez-Herrera A, Trueba M, Gomez-Muñoz A. 2020. Novel signaling aspects of ceramide 1-phosphate. BBA - Mol Cell Biol Lipids 1865:158630.

70. Maceyka M, Spiegel S. 2014. Sphingolipid metabolites in inflammatory disease. Nature 510:58–67.

71. Espaillat MP, Shamseddine AA, Adada MM, Hannun YA, Obeid LM. 2015. Ceramide and sphingosine-1-phosphate in cancer, two faces of the sphinx. Transl Cancer Res 4.

72. Granata R, Trovato L, Garbarino G, Taliano M, Ponti R, Sala G, Ghidoni R, Ghigo E. 2004. Dual effects of IGFBP-3 on endothelial cell apoptosis and survival: Involvement of the sphingolipid signaling pathways. FASEB J 18:1456–1458.

73. Varma Shrivastav S, Bhardwaj A, Pathak KA, Shrivastav A. 2020. Insulin-Like Growth Factor Binding Protein-3 (IGFBP-3): Unraveling the Role in Mediating IGF-Independent Effects Within the Cell. Front Cell Dev Biol 8:286.

74. Blázquez C, Carracedo A, Salazar M, Lorente M, Egia A, González-Feria L, Haro A, Velasco G, Guzmán M. 2008. Down-regulation of tissue inhibitor of metalloproteinases-1 in gliomas: a new marker of cannabinoid antitumoral activity? Neuropharmacology 54:235–243.

75. Rath GM, Schneider C, Dedieu S, Sartelet H, Morjani H, Martiny L, El Btaouri H. 2006. Thrombospondin-1 C-terminal-derived peptide protects thyroid cells from ceramide-induced apoptosis through the adenylyl cyclase pathway. Int J Biochem Cell Biol 38:2219–2228.

76. Yamanaka M, Shegogue D, Pei H, Bu S, Bielawska A, Bielawski J, Pettus B, Hannun YA, Obeid L, Trojanowska M. 2004. Sphingosine kinase 1 (SPHK1) is induced by transforming growth factor-β and mediates TIMP-1 up-regulation. J Biol Chem 279:53994–54001.

77. Kannan R, Jin M, Gamulescu M-A, Hinton DR. 2004. Ceramide-induced apoptosis: role of catalase and hepatocyte growth factor. Free Radic Biol Med 37:166–175.

78. Kulhankova K, Kinney KJ, Stach JM, Gourronc FA, Grumbach IM, Klingelhutz AJ, Salgado-Pabón W. 2018. The superantigen toxic shock syndrome toxin 1 alters human aortic endothelial cell function. Infect Immun 86:e00848–17.

79. Venter C, Niesler C. 2019. Rapid quantification of cellular proliferation and migration using ImageJ. Biotechniques 66:99–102.

80. Carpentier G. 2012. ImageJ Contribution: Angiogenesis Analyzer. ImageJ News 5.

81. Zhu W-H, Nicosia RF. 2002. The thin prep rat aortic ring assay: A modified method for the characterization of angiogenesis in whole mounts. Angiogenesis 5:81–86.

82. Stiffey-Wilusz J, Boice JA, Ronan J, Fletcher AM, Anderson MS. 2001. An *ex vivo* angiogenesis assay utilizing commercial porcine carotidartery: Modification of the rat aortic ring assay. Angiogenesis 4:3–9.

