## Supplemental Figures for "*Staphylococcus aureus* β-toxin exerts anti-angiogenic effects by inhibiting re-endothelialization and neovessel formation"

### Supplemental Figure 1

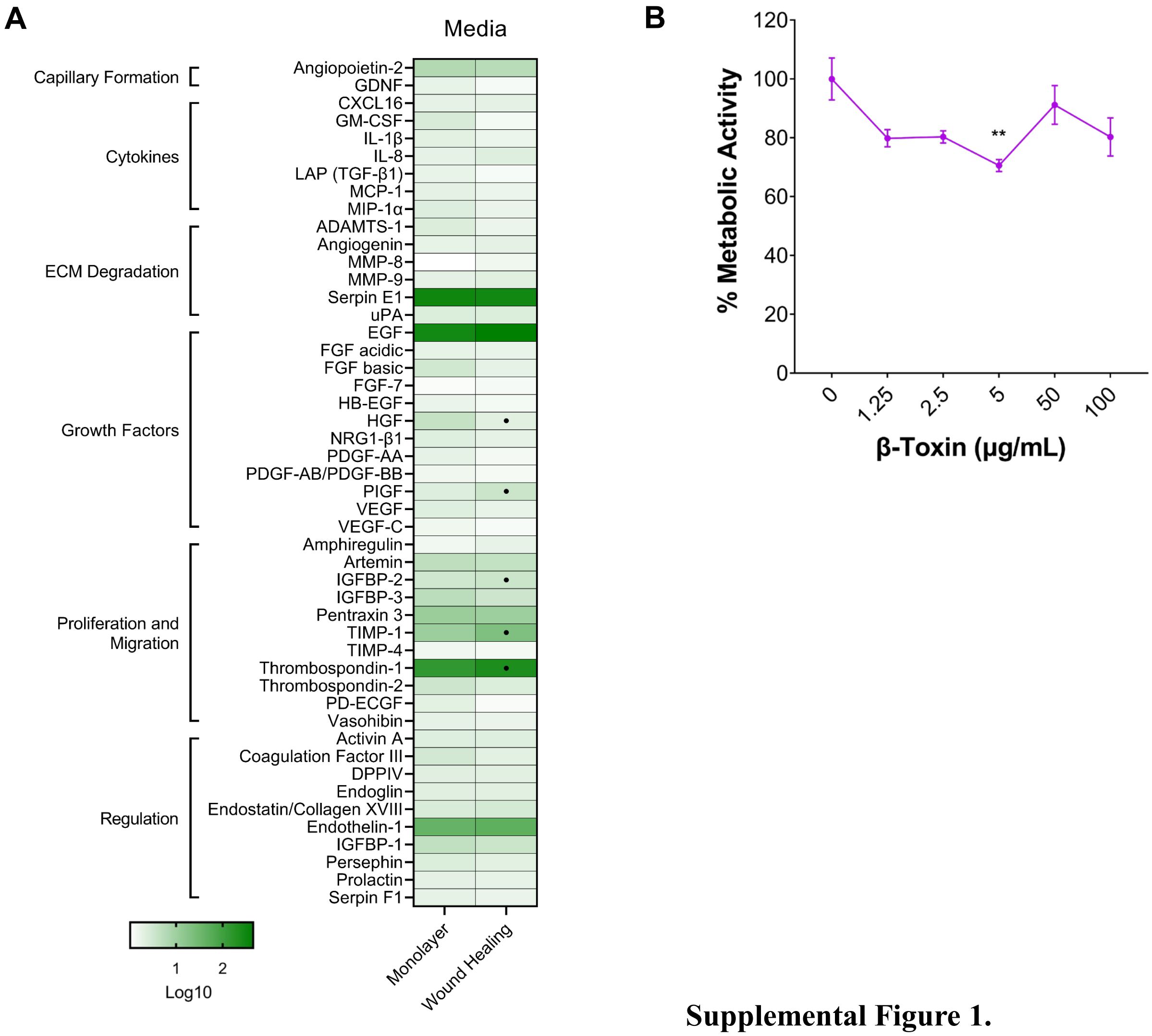


**Supplemental Figure 1.**

(A) Proteome analysis of untreated iHAECs grown to confluency on 1% gelatin-coated plates. Data is log scale with background threshold removed. • angiogenic-related factors with a 50% increase (>1.5-fold change) or decrease (<0.5-fold change) from media control.

(B) Percent metabolic activity. iHAECs were grown to near confluency on 1% gelatin-coated plates and treated for 24 h with β-toxin (50 µg $mL^{-1}$). Statistical significance determined by unpaired, two-tailed t-tests compared to untreated (0 µg $mL^{-1}$).

### Supplemental Figure 2

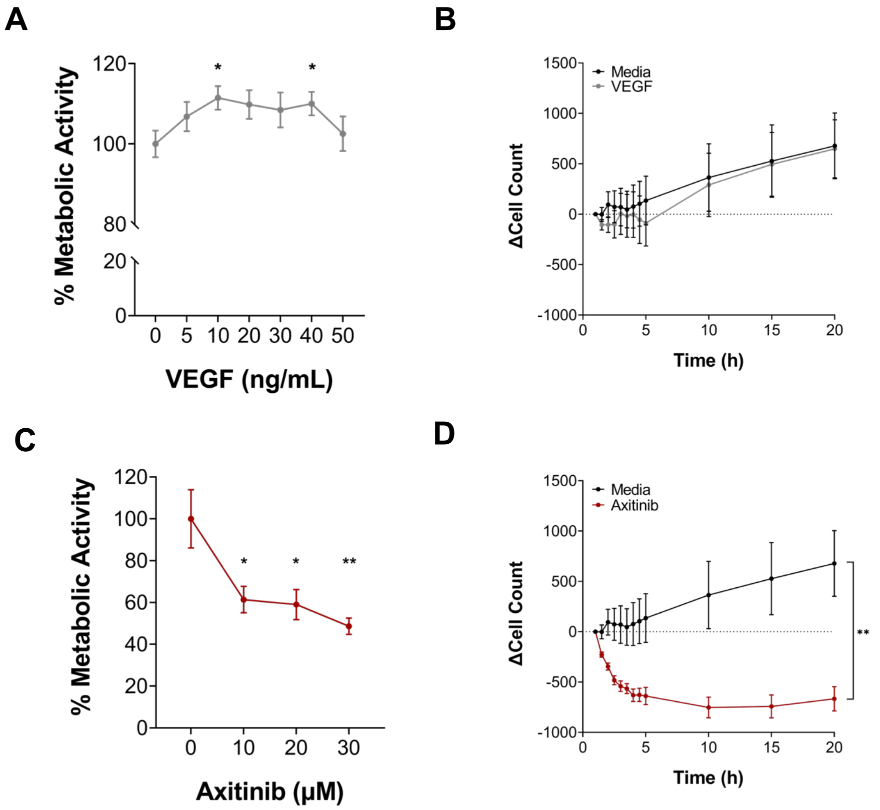


**Supplemental Figure 2.**

(A, C) Percent metabolic activity. iHAECs were grown to near confluency on 1% gelatin-coated plates and treated for 24 h with (A) VEGF or (C) axitinib. Statistical significance was determined by unpaired, two-tailed t-tests compared to untreated (0 µg $mL^{-1}$).

(B, D) Cell proliferation of iHAECs seeded at 7,000 cells $well^{-1}$ and treated with (B) VEGF (10 ng $mL^{-1}$) or (D) axitinib (10 µM) over a 20-h period. Cells counted every 30 min for the first 5 h then every 5 h thereafter. Results represent the change in cell count (mean ± SEM) of three independent experiments conducted in triplicate. Statistical significance determined by unpaired, two-tailed t-test at 20 h.

### Supplemental Figure 3

A

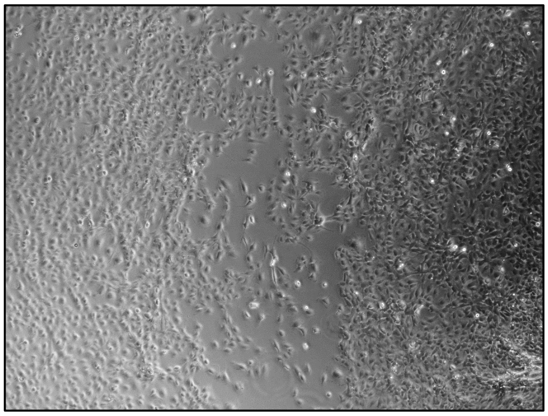

B

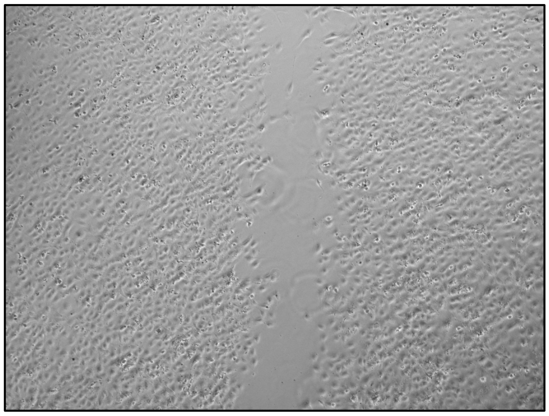

C

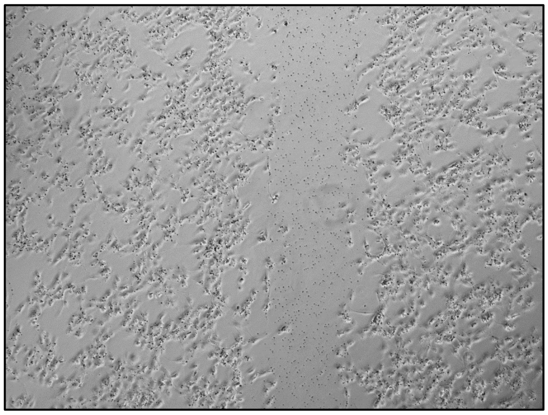

**Supplemental Figure 3.**

Phase-contrast microscopy of wound healing experiments at 24 h.

(A) Untreated iHAECs.

(B) iHAECs treated with 10 μM axitinib.

(C) iHAECs treated with 30 μM axitinib.

### Supplemental Figure 4

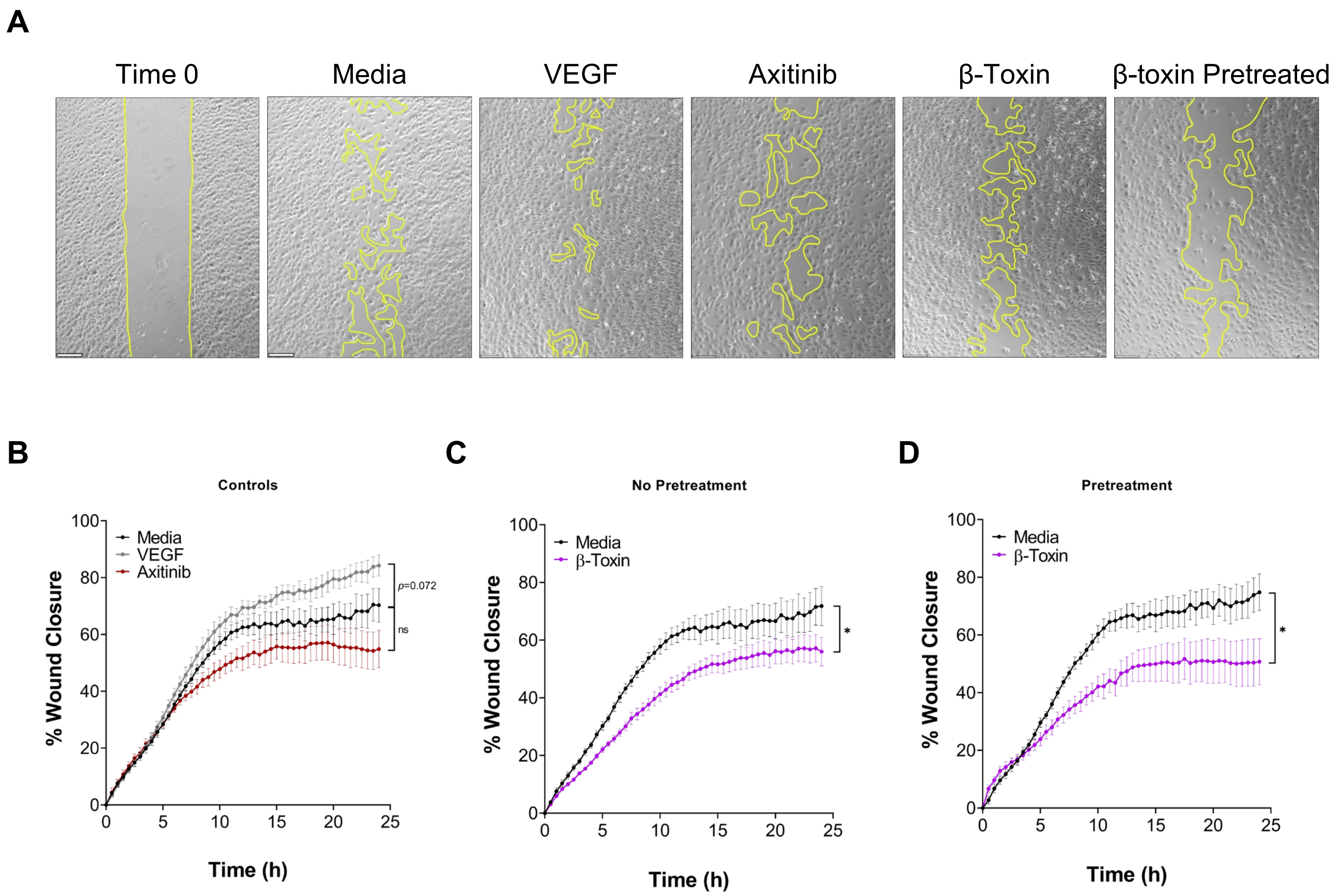


**Supplemental Figure 4.**

(A) Phase-contrast microscopy Time 0 (representative image) and at 24 h. Images captured every 30 min. Scale bar = 200 µm.

(B) HUVECs treated with VEGF (10 ng $mL^{-1}$) or axitinib (10 µM) at the start of the experiment.

(C) HUVECs treated with β-toxin (50 µg $mL^{-1}$) at the start of the experiment.

(D) HUVECs treated overnight with β-toxin (50 µg $mL^{-1}$) prior to gap formation and thereafter.

(B-D) All results are mean ± SEM of five independent experiments with four replicates each. * $p < 0.0332$; two-way repeated measures ANOVA.

### Supplemental Figure 5

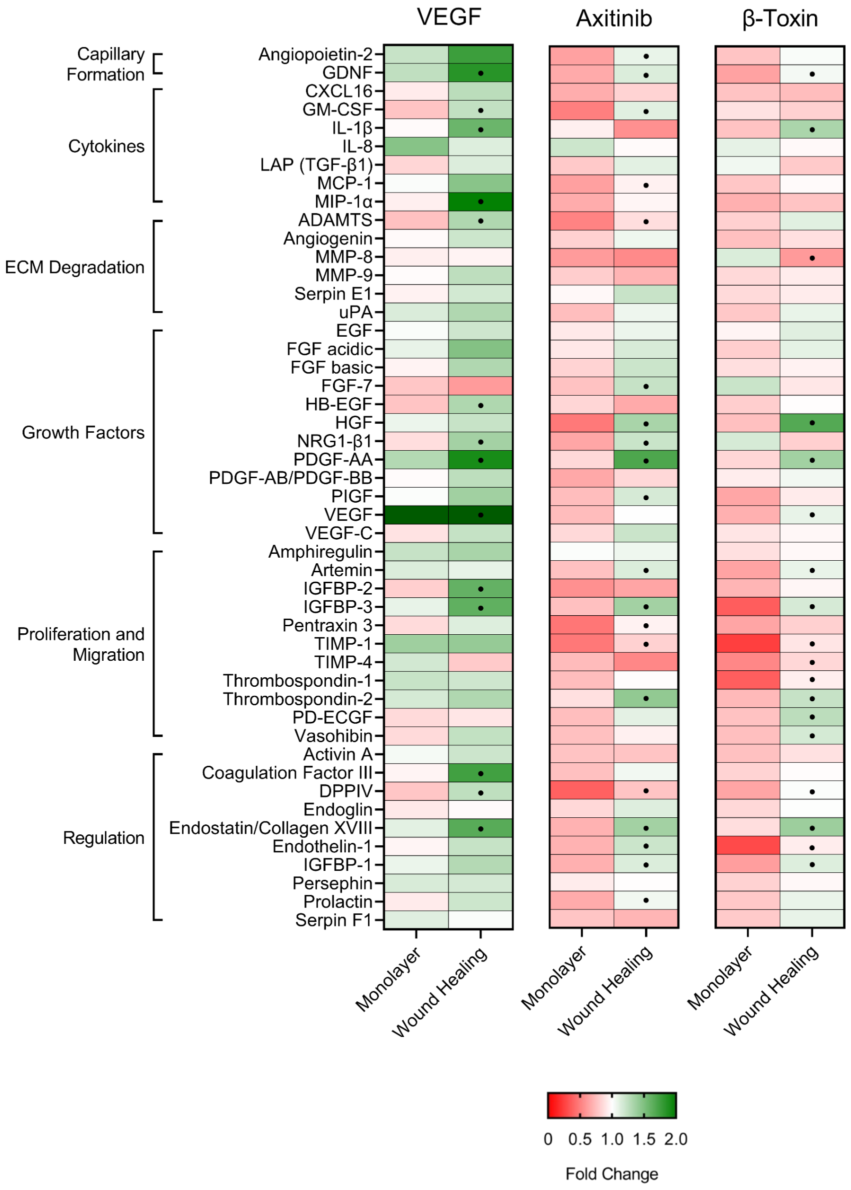


**Supplemental Figure 5.**

iHAECs treated with VEGF (10 ng mL$^{-1}$), axitinib (10 µM in wound healing; 30 µM in monolayers), or β-toxin (50 µg mL$^{-1}$). Data is on a linear scale and is a reproduction of data in Figures 1 and 3. Results are the mean fold change over matched untreated cells• angiogenic-related factors with a 50% increase (>1.5-fold change) or decrease (<0.5-fold change) from media control.

### Supplemental Figure 6

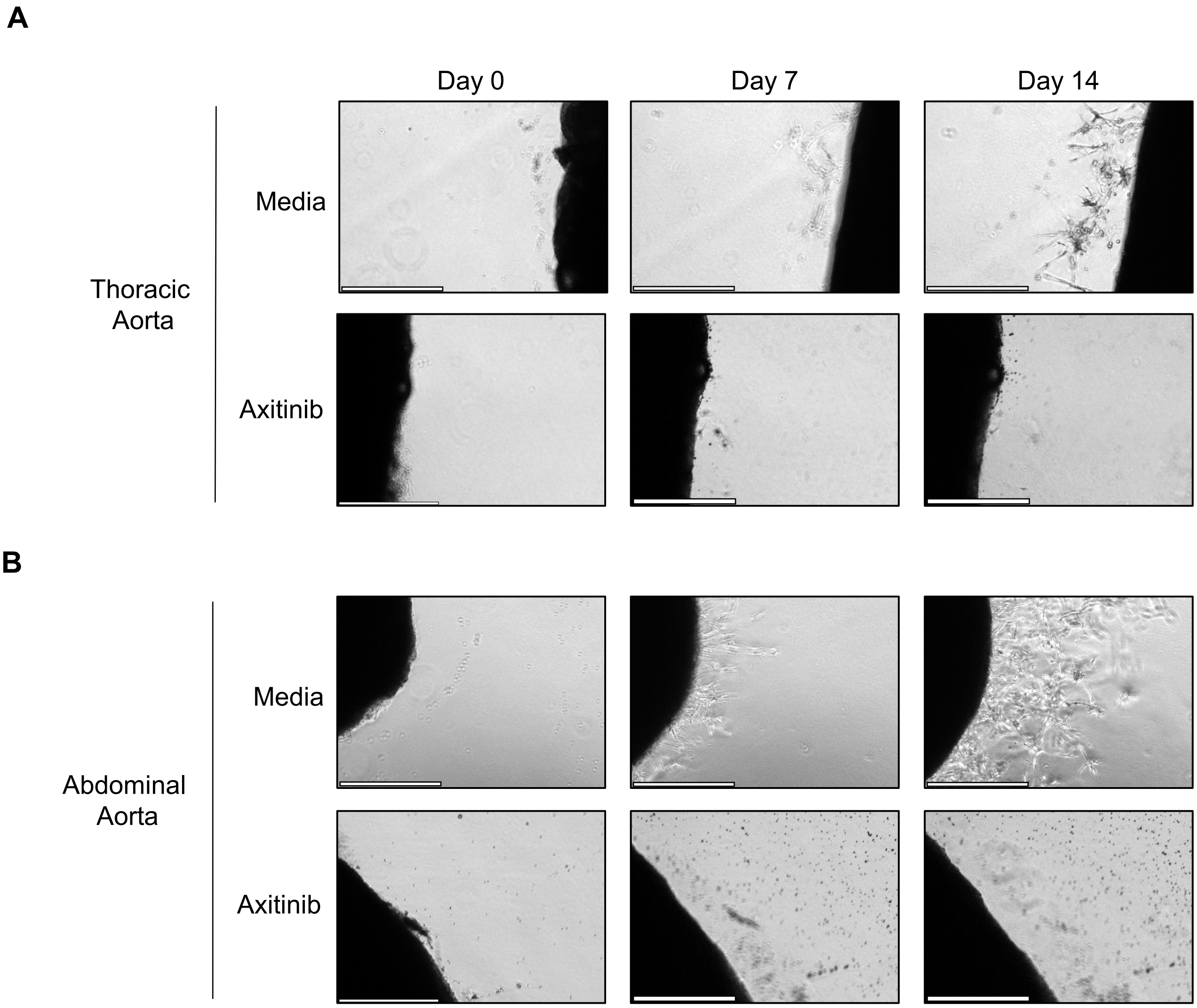


**Supplemental Figure 6.**

Thoracic and abdominal aortas were collected and sectioned from 2–3 kg New Zealand white rabbits. Rings were cultured on GFR-Matrigel in the presence or absence of axitinib (10 μM). Scale bar = 500 μm.

(A) Phase-contrast microscopy of thoracic aortic rings.

(B) Phase-contrast microscopy of abdominal aortic rings.

### Supplemental Figure 7

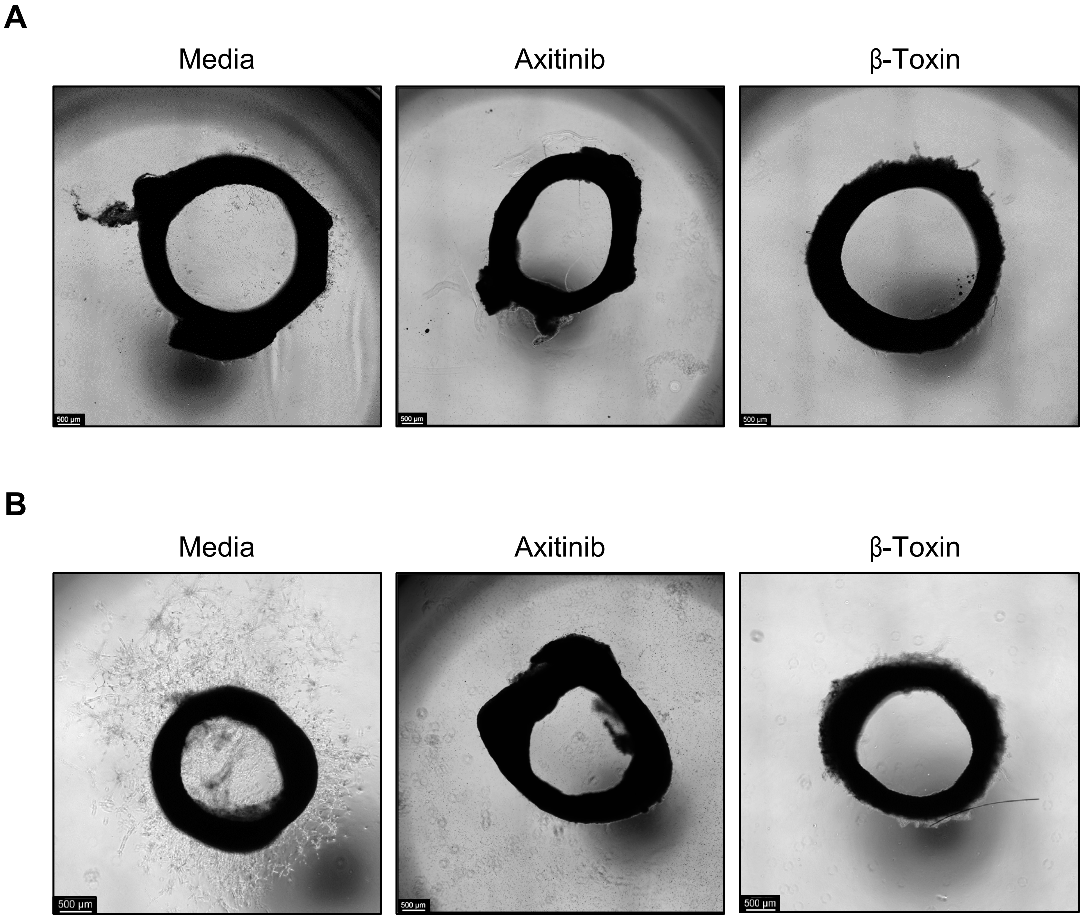


**Supplemental Figure 7.**

Thoracic and abdominal aortas were collected and sectioned from 2–3 kg New Zealand white rabbits. Rings were cultured on GFR-Matrigel in the presence or absence of axitinib (10 µM). Scale bar = 500 µm.

(A) Phase-contrast microscopy of thoracic aortic rings.

(B) Phase-contrast microscopy of abdominal aortic rings.
